# Spatial Capture-Recapture for Categorically Marked Populations with An Application to Genetic Capture-Recapture

**DOI:** 10.1101/265678

**Authors:** Ben C. Augustine, J. Andrew Royle, Sean M. Murphy, Richard B. Chandler, John J. Cox, Marcella J. Kelly

## Abstract

Recently introduced unmarked spatial capture-recapture (SCR), spatial mark-resight (SMR), and 2-flank spatial partial identity models (SPIM) extend the domain of SCR to populations or observation systems that do not always allow for individual identity to be determined with certainty. For example, some species do not have natural marks that can reliably produce individual identities from photographs, and some methods of observation produce partial identity samples as is the case with remote cameras that sometimes produce single flank photographs. These models share the feature that they probabilistically resolve the uncertainty in individual identity using the spatial location where samples were collected. Spatial location is informative of individual identity in spatially structured populations with home range sizes smaller than the extent of the trapping array because a latent identity sample is more likely to have been produced by an individual living near the trap where it was recorded than an individual living further away from the trap. Further, the level of information about individual identity that a spatial location contains is determined by two key ecological concepts, population density and home range size. The number of individuals that could have produced a latent or partial identity sample increases as density and home range size increase because more individual home ranges will overlap any given trap. We show this uncertainty can be quantified using a metric describing the expected magnitude of uncertainty in individual identity for any given population density and home range size, the Identity Diversity Index (IDI). We then show that the performance of latent and partial identity SCR models varies as a function of this index and produces imprecise and biased estimates in many high IDI scenarios when data are sparse. We then extend the unmarked SCR model to incorporate partially identifying covariates which reduce the level of uncertainty in individual identity, increasing the reliability and precision of density estimates, and allowing reliable density estimation in scenarios with higher IDI values and with more sparse data. We illustrate the performance of this “categorical SPIM” via simulations and by applying it to a black bear data set using microsatellite loci as categorical covariates, where we reproduce the full data set estimates with only slightly less precision using fewer loci than necessary for confident individual identification. The categorical SPIM offers an alternative to using probability of identity criteria for classifying genotypes as unique, shifting the “shadow effect”, where more than one individual in the population has the same genotype, from a source of bias to a source of uncertainty. We discuss the difficulties that real world data sets pose for latent identity SCR methods, most importantly, individual heterogeneity in detection function parameters, and argue that the addition of partial identity information reduces these concerns. We then discuss how the categorical SPIM can be applied to other wildlife sampling scenarios such as remote camera surveys, where natural or researcher-applied partial marks can be observed in photographs. Finally, we discuss how the categorical SPIM can be added to SMR, 2-flank SPIM, or other future latent identity SCR models.

## 1 Introduction

Animal population density is a fundamental concept in wildlife ecology and therefore, estimating population density is a primary challenge for ecologists (Efford, 2004; Laake *et al*., 1993). Mark-recapture and spatial mark-recapture (SCR) methods are among the most reliable methods for estimating population abundance and density; however, they generally require that individual identities of captured animals are determined with certainty (e.g., marks are not lost and recorded correctly Otis *et al*., 1978). Recently, several classes of spatial capture-recapture (SCR) that utilize latent or partially latent individual identities have been introduced, extending the utility of SCR models to populations that are unmarked (unmarked SCR; Chandler & Royle, 2013), populations for which only a subset of individuals are marked (spatial mark-resight (SMR); Sollmann *et al*., 2013), and populations for which some or all samples carry only partial identifications (spatial partial identity models (SPIM); Augustine *et al*., in press). All of these models share the feature that the true capture histories for some or all individuals that produced the observed samples are latent and must be probabilistically reconstructed by updating the latent individual identity of each sample in an MCMC algorithm. Further, all of these models use the spatial location where samples were collected, together with a spatially explicit model of sample deposition, to probabilistically link latent or partial identity samples collected closer together in space more often than those collected further apart, reducing the magnitude of uncertainty in individual identity and thus, population density estimates. Therefore, the spatial location where a sample was collected constitutes a continuous partial identity and unmarked SCR and SMR can be considered special cases of a SPIM.

The key feature of a SPIM is that the magnitude of uncertainty in individual identity and the model used to resolve it stems directly from two key aspects of population ecology-population density and home range size. In models with latent individual identities, there will be more uncertainty when more individuals are exposed to capture at the same traps, which occurs when animals live at higher densities and when their home ranges are larger. Within the context of an SCR model, this is equivalent to scenarios in which there is a higher density of individual activity centers and larger detection function spatial scale parameters, e.g., *σ* for the common half-normal detection model. Note, however, that the key feature that both population density and home range size determine which is directly responsible for the magnitude of uncertainty in individual identity is the magnitude of home range overlap, or more conceptually, the magnitude of overlap in individual utilization distributions (e.g., as quantified by Fieberg & Kochanny, 2005), which increases independently by increasing either population density or home range size.

We propose a metric to quantify the degree of home range overlap for a given population density and *σ*, and thus, the expected magnitude of uncertainty in individual identity. The Simpson’s Diversity Index can be applied to the individual detection probabilities averaged over many points on the landscape and realizations of the SCR process model to produce an Identity Diversity Index (IDI), which is conceptually similar to a metric of utilization distribution overlap for all individuals in the population. The details of how to calculate the proposed IDI can be found in Appendix A, but Figure 1 provides a visualization of how the magnitude of uncertainty in individual identity scales with population density and home range size as quantified by the SCR *σ* parameter. This relationship between the magnitude of uncertainty in individual identity and the spatial features of animal populations are what set a SPIM apart from the recently introduced non-spatial partial identity models (e.g. Bonner & Holmberg, 2013; McClintock *et al*., 2013; Knapp *et al*., 2009), where the magnitude of uncertainty in individual identity scales with population abundance alone.

Another key feature of the currently available SPIM models is that partial identity information can reduce the uncertainty in individual identity through three mechanisms-by adding deterministic identity associations, adding deterministic identity exclusions, and by improving probabilistic identity associations. Here, we define a deterministic identity association as a connection between samples from the same individual that also implies that the samples are excluded from being connected with samples from other individuals. This is distinguished from a deterministic identity exclusion, which can only prevent certain samples from being combined together. A probabilistic identity association occurs when two samples have a positive posterior probability of belonging to the same individual, and as this probability increases, the probability they belong to another individual necessarily decreases. Probabilistic identity associations can be improved with partial identity information, effectively converging to deterministic identity associations as partial identity information increases.

All SPIM models use the spatial location where samples were collected to improve probabilistic identity associations. The unmarked SCR (Chandler & Royle, 2013) uses this information alone to inform individual identity. Typical SCR, on the other hand, uses all possible deterministic identity associations. SMR (Sollmann *et al*., 2013) and the 2-flank SPIM (Augustine *et al*., in press) represent two intermediate cases that both utilize some deterministic identity associations and exclusions. SMR makes deterministic identity associations between the samples of identifiable individuals, usually marked, but potentially unmarked, which simultaneously excludes them from being connected to samples from other individuals. Deterministic identity exclusions are then made between the samples of unidentifiable individuals whose mark status can be observed (e.g., an unmarked sample cannot belong to a marked individual Royle *et al,* 2013). The 2-flank SPIM makes deterministic identity associations across the same flank of the same individual, enforcing exclusions with non-matching samples from the same flank. Deterministic identity exclusions arise in the 2-flank SPIM from the fact that an individual can only have one left and right flank.

Augustine *et al.* (in press) demonstrated that further deterministic identity exclusions are possible in SPIMs by using individual sex to split a data set into two population groups whose latent identities could not logically match, reducing the uncertainty in individual identity and thus abundance and density estimates. Splitting the population into identity subgroups of increasingly smaller size is conceptually similar to applying unmarked SCR to separate populations, each with increasingly lower population densities, moving the population under study to more favorable regions of the IDI along the density dimension (Figure 1). However, rather than splitting data sets into increasingly small subsets, it is desirable to have a model that incorporates these categorical identity exclusions, allows for imperfect observation of the category levels, and allows parameters to be shared across population identity subgroups. Further, when all categories are combined into a single analysis, the distribution of individuals across the category levels provides further information that can improve the probabilistic identity associations. For example, if a population is 75% female, it is more likely that two nearby male samples came from a single individual than if the population is 75% male. Thus, we are introducing a new class of SCR model, the “categorical SPIM”, which uses partially identifying identity covariates to add both deterministic identity exclusions and reduce the uncertainty in probabilistic identity associations.

Partially-identifying categorical covariates exist in many types of invasive and noninvasive wildlife sampling; for example, in studies using remote cameras, features such as sex, age class, and color morph may be observable in at least some photographs. In more invasive wildlife sampling involving live capture, many more features are measurable and researchers may apply categorical marks whose combination do not provide full identities (e.g., colored collars or ear tags), or categorical marks may be fully identifying (e.g., Lewis *et al.*, 2015), but imperfectly observed. Perhaps the most informative source of categorical identity covariates come from microsatellite genotypes, which we will use as the main application to demonstrate how the categorical and spatial information combine to determine the magnitude of uncertainty in individual identity.

In genetic mark-recapture, individual identities are constructed from amplified microsatellite loci – non-coding, highly variable segments of the genome-with individuals considered individually unique when a sufficient number of loci are amplified that it is very unlikely that any two individuals in the population share the same multilocus genotype. The number of loci necessary to guarantee uniqueness of all genotypes in a population with near certainty depends on the diversity of genotypes and the genotype frequencies at each microsatellite loci (Waits *et al.*, 2001). This determination is typically made using P(ID) and/or P(sib) criteria, both of which estimate the probability that any two randomly selected individuals (or full siblings) in a population would share the same multilocus genotype given the observed allele diversities and frequencies (Waits *et al.*, 2001).

The standard practice in genetic capture-recapture is to use enough microsatellite loci such that P(ID) or P(sib) are less than a strict threshold, such as 0.05 or 0.01. Although this approach can effectively minimize the number of individuals erroneously classified as the same individual in a capture-recapture data set, these errors cannot be completely eliminated with absolute certainty. The possibility that multiple individuals have the same multilocus genotype in a capture-recapture data set has been referred to as the “shadow effect” (Mills *et al.*, 2000) and is considered a type of low-frequency error in assigning individual identities that introduces minimal bias into parameter estimates in capture-recapture studies as long as P(ID) or P(sib) criteria are strictly enforced. Using the categorical SPIM to model genotype data is especially appealing as it does not make deterministic connections between samples with matching genotypes and thus, multiple individuals in the population may have the same genotype, shifting the “shadow effect” from a source of bias to an additional source of uncertainty.

Here, we generalize the unmarked SCR model to develop the “categorical SPIM”. We show via simulation that in scenarios with more sparse data than previously considered and/or scenarios with larger *sigmas* and larger densities, the unmarked SCR density estimator is biased and very imprecise, demonstrating the importance of population density and home range size to the application of latent and partial identity SCR models. We then show that adding categorical identity covariates removes this bias and increases precision, allowing for reliable density estimation across a wider range of values of density and *σ* for a given capture process scenario. We also demonstrate that the uncertainty in the posterior for *n^cap^,* the latent number of individuals captured during a survey, correlates well with the uncertainty in the posterior of *N,* suggesting it is a good single metric to quantify the observed magnitude of uncertainty in individual identity for a given survey. Finally, we apply the categorical SPIM to a previously-published black bear data set in which we demonstrate how well the proposed model can reproduce the original density estimate using fewer loci than originally genotyped. Using this data set, we demonstrate that all uncertainty in individual identity can be removed with enough categorical identity covariates, producing equivalent estimates to an SCR model where all identities are known with certainty.

## 2 Methods – Data and Model Description

### 2.1 Methods – Unmarked SCR Foundation

First, we introduce the version of the unmarked SCR model that we will expand to allow categorical identity covariates. The unmarked SCR model is a typical hierarchical SCR model except that information about individual identity is not retained during the observation process. Formal inference is achieved by relating the spatial pattern of observed counts or detections at each of the *J* traps to the latent structure of the SCR process model. For the process model, we assume the *N* individuals in the population have 2-dimensional activity centers that are distributed uniformly across a two-dimensional state space *S* of arbitrary size (*A*) and shape, i.e. *s_i_* ~ Uniform(***S***), *i* = 1, …, *N* (see Borchers & Efford, 2008; Reich & Gardner, 2014; Royle *et al.*, 2016, for alternative specifications). The activity centers are organized in the *N* × 2 matrix S.

For the observation model, we introduce the *N* × *J* fully latent capture history ***Y**^true^*, recording the number of detections or counts for each individual at each trap summed across the *K* occasions. The locations of the *J* traps are stored in the *J* × 2 matrix *X*. We assume that the number of counts or detections for each individual at each trap is a decreasing function of distance between the activity centers and traps. If using a count model, we assume the latent counts are Poisson: 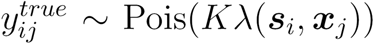, where 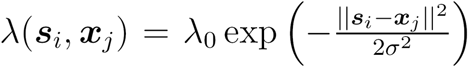, *x_j_* is the location of trap *j*, *λ*_0_ is the expected number of counts for a trap located at distance 0 from an activity center, and *σ* is the spatial scale parameter determining how quickly the expected counts decline with distance. We also consider an alternative Bernoulli observation model for which 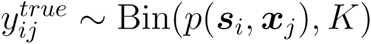, where *p*(*s_i_*, *x_j_*) = 1 – exp(–λ(*s_i_*, *x_j_*)).

During the observation process, the true, latent capture history, ***Y**^true^*, is disaggregated into the observed capture history, ***Y***^obs^, discarding information about individual identity by storing one observation per row in ***Y**^obs^* (e.g., no samples are deterministically connected to the same individual). More specifically, ***Y**^obs^* is the *n^obs^* × *J* matrix with entries 1 if sample *m* was recorded in trap *j* and 0 otherwise. Note that if we assume a Bernoulli observation model, each detection event will constitute a single observation, while if we assume a Poisson observation model, counts are disaggregated into observations of single counts, because counts from the same individuals cannot be deterministically connected without certain and unique identities. To visualize this, below is an example of true and disaggregated observed data set where *N*=2 and *J=*3:

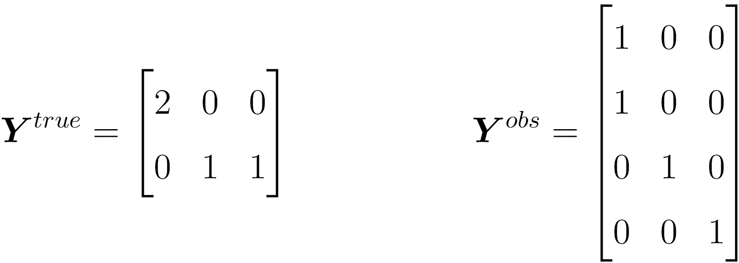

### 2.2 Methods – Categorical SPIM

We propose a class-structured version of the unmarked SCR model in which class membership is determined by each individual’s full categorical identity (e.g., full genotype). Here, we define a *full categorical identity* to be an individual’s set of true values for *n_cat_* categorical covariates, where *n_cat_* is the maximum number of categorical covariates considered, and multiple individuals in the population can share the same full categorical identity. We will modify this model such that single or multiple categorical covariates, potentially partially or even fully unobserved, are recorded with each observed sample at each trap. Further, continuous covariates could be discretized into categories if it is safe to assume there is no measurement error. The linked density and categorical covariate models (joint process model) are fully latent and we use the *n^obs^* trap-referenced, observed categorical covariates to make inference about this latent structure. The observed data then consist of two linked data structures: ***Y***^obs^, an *n^obs^* × *J* capture history indicating the trap at which each sample was recorded and ***G***^obs^, an *n^obs^* × *n^cat^* identity history recording the observed categorical covariate(s) of each sample with category level enumerated sequentially as described below, or recorded as a 0 if not observed.

For the joint process model, we assume that each individual has a full categorical identity associated with its activity center. Following Wright *et al*. (2009), we assume that all possible category levels for each categorical covariate are known, with the number of categories for each covariate *l* being 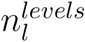, *l* = 1,…, *n^cat^*. Next, we introduce the population category level probabilities for category *l*, *γ*_*l*_, of length 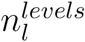 and corresponding to the enumerated category levels (1,…, 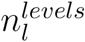) for covariate *l*. Then we introduce the *N* × *n^cat^* matrix ***G**^true^*, where 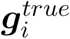 is the full categorical identity of the individual with activity center *S_i_*. Finally, we assume the categorical identity of each individual for each covariate are distributed following the covariate-specific category level probabilities according to 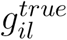 ∼ Categorical(*γ_l_*), implying that category levels are independent across covariates (e.g., linkage equilibrium in the genetic context) and individuals. Using the example true and observed capture histories above, potential true and observed structures for the categorical identities are:

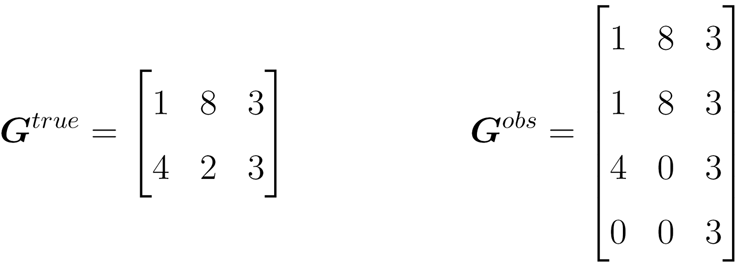

In this case, the first two observed samples could possibly have come from the same individual as could the third and fourth sample; however, the third sample could not have come from the same individual as the first two samples. The fourth sample with two unobserved categories could possibly belong to the same individual as that which produced the first three samples, with only the third sample being a correct match.

The observation process is the same as unmarked SCR except that a categorical identity, potentially partially or fully latent, is associated with each trap-referenced observation. The missing data process could be a simple binomial model for identification success, perhaps with covariate-specific identification probabilities; however, if we assume that covariate observation values do not vary by individual or by category level, the likelihood for the missing data process does not change when updating latent identities or latent categorical covariate values and can be ignored in the MCMC algorithm. Therefore, we will make the assumption that covariate identification probabilities do not vary by individual or category level; however, this assumption would be easy to relax.

The unmarked SCR model and categorical SPIM use a process similar to data augmentation to estimate population abundance and density (Royle *et al*., 2007) and to model the uncertainty in individual identity by providing latent structure to allow for different configurations of the observed samples across the individuals in the population (Chandler & Royle, 2013; Augustine *et al*., in press). Unlike typical data augmentation, the number of captured individuals, *n^cap^*, in unmarked SCR is unknown, so rather than augmenting the observed capture history, a fully latent capture history, ***Y**^true^*, of size *M* × *J* is defined and ***Y**^true^* is initialized by assigning the observed samples in ***Y**^obs^* an individual identity based on the spatial proximity of samples and the compatiblity of their observed categorical identities.

We then augment ***G**^true^* to size *M* × *n^cat^*, which is initialized using the minimally-implied categorical identity of the samples each individual is initialized with and the remaining elements of ***G**^true^* that are not determined by these samples are simulated from the category level probabilities. Similar to typical data augmentation, *M* is chosen by the analyst to be much larger than *N* and we use **z,** a latent indicator vector of length *M*, to indicate which individuals are in the population; however, unlike typical data augmentation, this vector is fully, rather than partially latent. We assume *z_i_* ~ Bernoulli(*ϕ*), inducing the relationship *N* ~ Binomial(M, *ϕ*). Then population abundance is 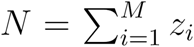 and population density, *D*, is *N/A*. See Appendix B for a full description of the MCMC algorithm.

Note that *n^cap^*, the total number of individuals captured, typically denoted by *n* and a known statistic in capture-recapture models, is a derived, random variable in unmarked SCR with a posterior distribution that quantifies the magnitude of uncertainty in individual identity. As more categorical identity information is added, the posterior distribution of *n^cap^* should converge to the single, true value. Finally, we introduce the derived vector, *I*, of size *n^obs^*, that records the latent individual, 1,…, *M*, each sample is assigned to. This vector is updated on each MCMC iteration, producing a posterior for true identity for each sample which can be post-processed to obtain pairwise posterior probabilities that any two samples originated from the same individual. The posterior distribution of the true covariate values of samples with missing values can also be recorded.

## 3 Simulations

### 3.1 Simulation Specifications

We conducted two simulation studies to explore the performance of the categorical SPIM. First, we conducted a simulation study to demonstrate that adding an increasing number of identity categories removes bias from the unmarked SCR density estimator and increases accuracy and precision. Here, the number of identity categories is defined to be the total number of unique categories implied by the *n_cat_* covariates with 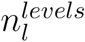 each. These simulations were also constructed to demonstrate that the performance of unmarked SCR and the effectiveness of adding identity categories varies as a function of population abundance, density given abundance, and *σ*.

We start with the sampling scenario similar to that of Chandler & Royle (2013); however, we make two modifications to create scenarios more challenging for the unmarked SCR estimator. Chandler & Royle (2013) explored a range of optimistic sampling scenarios as a proof of concept for the unmarked SCR estimator, where sampling takes place on a 15 x 15 trapping array with unit spacing within a 20 x 20 state space, *λ*_0_ = 0.5, *σ* ∈ {0.5,0.75,1}, and *D* ∈ {0.0675,0.1125,0.1875}, corresponding to *N* ∈ {27,45,75}. We consider scenarios with higher *D* given *N*, achieved by decreasing the size of the trapping array and state space, and with more sparse detection data. We increased data sparsity by reducing *λ*_0_ to 0.25 for the first two scenarios and further for the subsequent two scenarios (see below), and reducing the number of traps by 64% from 225 to 81 (9 x 9 grid with unit spacing, buffered by 3 units). We considered higher densities for similar values of *N* by specifying *D* ∈ {0.17,0.35}, corresponding to *N* ∈ {39,78} the latter *D* being larger than explored by Chandler & Royle (2013) for a similar *N* (75). We considered that the populations were sampled for *K =* 10 occasions (Chandler & Royle, 2013, considered *K* ∈ (5,10)).

We conducted simulations across 4 scenarios with a 2 x 2 factorial design using low and high values of *σ* and *D*. Scenarios A1 and A2 were the low *σ* scenarios with *σ* = 0.5, and Scenarios A3 and A4 doubled *σ* to 1. To account for compensation in the detection function parameters (Efford & Mowat, 2014) and maintain similar levels of data sparsity with the larger *σ*, we lowered *λ*_0_ from 0.25 to 0.061 to approximately match the expected number of captures for each individual to that of the scenarios with *σ* = 0.5 (E[caps]≈ 1.65, achieved by trial and error). On this unit spacing grid, with *σ* = 0.5, the majority of an individual’s captures fell within a 4-trap area, whereas with *σ* = 1 the majority of an individual’s captures fell within a 16-trap area. Scenarios A1 and A3 were the high abundance scenarios with N=78, and Scenarios A2 and A4 were the low abundance scenarios with N=39. The approximate Identity Diversity Indices (interpolated from Figure 1) for scenarios A1-A4 were 0.38, 0.23, 0.76, 0.58. Within each scenario, we explored 9 subscenarios with differing numbers of identity categories, with 1 identity category corresponding to unmarked SCR. Following the unmarked SCR subscenario, we sequentially added identity covariates with 2 category levels each, leading to the number of unique identity categories increasing exponentially with base 2 (2, 4, 8, 16, 32, 64, 128, 256).

Further, scenario A2 was modified to better disentangle the effects of increasing *D* from increasing *N*. Increasing *D* by increasing *N* simultaneously increases uncertainty in individual identity and reduces data sparsity, which have opposite effects on estimator precision. Therefore, to better explore the effect of increasing *D* has on the uncertainty in individual identity, *D* must be increased by constraining a fixed *N* into a smaller state space. The state space can be reduced in two ways; the number of traps can be reduced, keeping the same state space buffer, or the number of traps can be fixed, reducing the state space buffer. In the first scenario, data sparsity is increased since a lower proportion of individuals will be located on the interior of the trapping array and we suspect the absolute number of traps is important for unmarked SCR density estimation. In the second scenario, data sparsity is decreased because a larger proportion of individuals live on the interior of the trapping array. In order to retain the same number of traps, we chose to increased the density of scenario A2 by reducing the state space buffer from 3 to 1 units, thus constrainting the *N* individuals into a reduced state space area (Scenario A2b). The reduction in the state space area increased *D* from 0.17 to 0.32, and raising the approximate IDI value from 0.23 to 0.37.

We conducted a second simulation study to demonstrate that the categorical SPIM can accommodate partially-observed categorical identities and provide a proof of concept for using partial genotypes that are the result of failed DNA amplification, rather than as part of the study design as would be the case in the first set of simulations if identity categories were genotype loci. We used the parameter values from scenario A3 above, but introduced imperfect detection to the observed genotypes. We simulated data sets with 7 categorical identity covariates, each with 5 equally common category levels, and the category value for each categorical covariate was then observed with probability 0.5, leading to the average categorical identity being observed at 3.5 of the categorical identity covariates. We fit the categorical SPIM model to these data sets, assuming all partial categorical identities were usable (Scenario B1) or 75% of the partial categorical identities were usable as might be the case when using partial genotypes if a subset was deemed to be unreliable due to the likelihood of containing genotyping errors (Scenario B2). We then fit the null SCR model to the perfectly observed data for comparison (Scenario B3).

For simulation scenarios A, we simulated and fit our model to 144 data sets within each subscenario, and for simulation scenarios B, we simulated and fit our model to 128 data sets (due to cluster computing availability). We ran single MCMC chains starting from the simulated parameter values for 60,000 iterations, enough to produce effective sample sizes for abundance of 400 or more for the categorical SPIM scenarios with 2 or more identity categories. The mixing for most unmarked SCR scenarios with 1 identity category was very poor for these challenging scenarios and frequentist performance was unlikely to improve with longer chains. We calculated point estimates using the posterior mode and interval estimates using the highest posterior density (HPD) interval. We were interested in the frequentist bias and coverage of the categorical SPIM estimator, the accuracy of the estimator depicted visually by the variance and right skew of the sampling distribution and quantitatively by the mean squared error and coefficient of variation (100×posterior sd/posterior mode), and the precision, quantified by the mean 95% HPD interval width. Also of interest was the use of the precision of *n^cap^*, also quantified by the mean 95% HPD interval width, as a metric of uncertainty in the individual identity of observed samples that can predict the uncertainty in *N*.

### 3.2 Simulation Results

The unmarked SCR abundance estimator was right-skewed with high variance (Figures 2 and 3), except in Scenario A2 where both *σ* was small and abundance was low. The unmarked SCR estimator had a large mean 95% CI width relative to abundance in all scenarios. The unmarked SCR abundance estimates were negatively biased (Table B1; Appendix B) in the higher abundance scenarios (A1 and A3) and positively biased in the lower abundance scenario with the larger *σ* (A4) or larger density (A2b). Adding and increasing the number of identity categories reduced bias and increased precision in all scenarios; however, with diminishing returns as more identity categories were added. The reduction in mean 95% CI width for *n^cap^* by the introduction of identity categories was closely related to the reduction in mean 95% CI width for abundance; however, the relationship was not linear and varied by scenario (Figure 4). More identity categories were required to reach maximum precision when abundance was higher and *σ* larger. The largest improvement in precision and abundance with the addition of identity categories was seen in the low abundance, high density, low *σ* scenario (A2b), where the majority of uncertainty in abundance was removed with the addition of 1 2-level categorical covariate. Note that the precision of A2b converged to a lower value than scenario A2 because the same *N* = 39 individuals were constrained to be located within 1 unit of the trapping array, rather than 3 units, decreasing data sparsity.

Adding just a few identity categories changed the negative bias in abundance for the high abundance scenarios to positive bias, and magnified the positive bias in the low abundance scenarios, a pattern that was more pronounced in the large *σ* scenarios and which we discussed further in the application. However, this positive bias was removed by the addition of more identity categories. Positive bias in 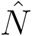 was <5% in Scenarios A1, A2, and A4 when 4 identity categories were available, and <5% in Scenario A3 when 16 identity categories were available. With no or very few identity categories, the latent identity samples tended to be allocated to more individuals than were actually captured, resulting in positive bias in *n^cap^*, with more bias in the large *σ* scenarios, and we attribute this as one cause for the positive bias in 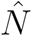.

Increasing *σ* decreased the precision and accuracy of the unmarked SCR and categorical SPIM *n^cap^* and *N* estimates (A1 vs. A3 and A2 vs. A4; Appendix B). Increasing *D* by increasing *N* decreased the precision and accuracy of the *n^cap^* estimates, but increased the precision and accuracy of the *N* estimates demonstrating that the additional uncertainty in individual identity was more than offset by the lower data sparsity (A1 vs. A2 and A3 vs A4). Increasing *D* without increasing *N* decreased the precision and accuracy of both *n^cap^* and *N* estimates. The IDI did not perfectly predict the metrics of precision and accuracy across scenarios that changed both *σ* and *D* for all levels of identity categories, but the IDI generally correlated negatively with precision and accuracy (Table B1).

The partially observed categorical identity covariate simulations produced minimally biased abundance point estimates (1 and 2.5% positive bias) and interval estimates that were nearly as precise as the scenario in which the identities were perfectly observed (Table 1). In these scenarios, categorical identities were observed at half of the 7 identity covariates, on average, producing data sets with an average of less than 1 sample observed at all category levels-data sets that would be unusable if full categorical identities were required, as might be the case with genetic capture-recapture. In Scenario B1, where all partial categorical identities were used, the interval estimate was 95% as precise as the complete data analysis, and in Scenario B2, where 25% of the partial categorical identities were unusable, the interval estimate was 80% as precise than the complete data analysis. In both scenarios, an average of 3.5 categorical covariates provided enough information that the uncertainty in *n^cap^* was small (mean credible interval widths of 1.8 and 2.4 relative to mean *n^cap^* of 18.5 and 17.0 in B1 and B2, respectively).

## 4 Application – Central Appalachian Black Bears

We applied the categorical SPIM model to an American black bear noninvasive hair trapping data set that used 7 microsatellite loci for individual identification. This data set comes from a study conducted along the Kentucky-Virginia, USA border across 2 study areas during 2012 and 2013 to estimate the population density and abundance of a recently reintroduced population that was in the process of recolonizing vacant range (Murphy *et al*., 2016). We chose to use the data set from the larger study area in 2013 because our model should perform better on the larger trapping array and more samples were collected in 2013 than in 2012 at this site. The specifics of the data collection methods are described by Murphy *et al*. (2016); of particular relevance is that eighty-one hair traps were deployed across the 215-km^2^ study area with an average trap spacing of 1.6 km, and all traps were checked weekly for 8 consecutive weeks, with a week constituting a capture occasion. Similar to most bear hair trapping studies, hair samples were subsampled for genotyping because of the prohibitive costs of genotyping thousands of samples, such that at most 1 hair sample per trap per occasion produced an individual identity. The capture and subsampling processes resulted in 95 samples from 45 females and 87 samples from 37 males, determined using the P(sib) criterion. The spatial distribution of traps and individually-identified hair sample observations are depicted in Figure 5. The microsatellites used were G10H, G10L, G10M, MU23, G10J, G10B, and G10P, which had genotype frequencies of 19, 22, 19, 17, 12, 15, and 10 for females and 21, 18, 18, 22, 14, 13, and 10 for males. Despite the large number of genotypes at each locus, the majority of individuals shared just 2-4 genotypes at each locus, making them less informative than if the loci-specific genotypes were equally distributed as they were in our simulation studies.

The goal of this analysis was to fit the categorical SPIM using from 1 to 7 loci, added in the order listed above, and to compare the estimates to the null SCR estimate that does not allow for any uncertainty in individual identity. Further, we also consider a scenario adding partial genotype samples (2 for females, 4 for males) into the analysis that were originally discarded. For all genotype scenarios, the trapping array was buffered in the X and Y dimension by 3 km for females and 6 km for males, leading to state space sizes of 1042.5 km^2^ and 1473.4 km^2^ for females and males, respectively. For each sex-specific, 1-7 loci data set, we ran 32 Markov chains for 250,000 iterations each, thinned by 50, and discarded the first 25,000 iterations as burn in, leaving 1.4 million samples from the posterior. This large number of posterior samples was likely far more than necessary; however, it allowed us to explore the behavior of the MCMC chains as the last uncertainty in *n^cap^* was removed by adding genotype information (see Supplement 1). Because the hair sample subsampling process allowed for at most 1 sample per individual/trap/occasion, we used a Bernoulli observation model. The metrics of comparison were the point estimates (posterior modes), posterior standard deviations, and coefficients of variation (100×posterior sd/posterior mode) for abundance as well as the posterior distributions of *n^cap^*. Note, however, that the analyses adding the partial genotypes should not be expected to reproduce the null SCR point estimates, standard deviations, and number of individuals captured because they included additional data not used by the null SCR estimate.

### 4.1 Application – Results

The categorical SPIM estimates (Table 2) for both sexes generally demonstrated the same patterns seen in the simulations. Abundance estimates with few identity categories were positively biased (relative to the SCR estimate) because the estimates of *σ* were negatively biased and/or the estimates of *n^cap^* were positively biased. The magnitude of bias was larger for males, either partially or fully the result of a larger *σ*, estimated to be 2 times larger than females; although. As more loci were added, the categorical SPIM abundance estimates and their posterior standard deviations converged towards those of the SCR model and the posterior modes of *n^cap^* converged towards the true number captured and the posterior variance of *n^cap^* converged towards 0 (Figure 6). The coefficient of variation for all categorical SPIM estimates was lower than 0.20, except for the 1 loci male estimate.

The 1 locus female estimate was only 30% percent less precise than the full SCR estimate, with a positive bias (relative to the complete data estimate) of 7.5%. The 3 loci female estimate was substantially better-11.5% less precise than the full SCR estimate and positively biased by only 2.5%. The 4 loci female estimate was effectively equivalent to the full SCR estimate, with 3% less precision and minimal bias. The 5-7 loci female estimates were negligibly improved. Adding 2 partial genotype samples reduced the posterior standard deviation by 1.7% and coefficient of variation less than 1%. One partial genotype sample was consistent with two different individuals in the full genotype data set, matching one with posterior probability of 0.452 and the other with posterior probability 0.544, leaving just a 0.004 posterior probability that this sample was from a new individual. The other partial genotype matched an individual with 2 captures in the full genotype data set with posterior probability 0.997, leaving a probability of 0.003 that this was a new individual, and adding a high probability spatial recapture for this individual.

The 1 locus male estimate was too biased (relative to the full SCR analysis) and imprecise to be of use, whereas the 2 and 3 loci male estimates had reasonable precision but perhaps too much positive bias to be useful. The 4 loci male estimate was 17% less precise than the full SCR estimate, with a positive bias of 7.7%. The 5 loci estimate is only negligibly less precise than the full SCR estimate, and adding the *6^th^* and *7^th^* loci improved precision negligibly. Adding 4 partial genotype samples modestly increased the abundance estimate, increased the posterior standard deviation (due to the larger point estimate), and decreased the coefficient of variation by 2.5%. The posterior probability that three of the partial genotype samples each came from separate individuals not represented in the full genotype data set was 1, while the fourth partial genotype sample matched with 8 other samples from 1 individual, each with posterior probability 1.

### 4.2 Application – Discussion

This analysis demonstrates that the categorical SPIM estimator performed similarly on a real world data set as it did for simulated data; however, the positive bias in the male estimates with few loci was larger than seen in the simulated data sets. We suspect individual heterogeneity in detection function parameters, particularly *σ*, may have been present in the male bear data set. If so, this could have led to poorer performance with few loci/identity categories, and the requirement of more loci/identity categories to remove bias and increase precision than if there were no individual heterogeneity. The distribution of observed spatial recaptures in Figure 5 does seem to suggest individual heterogeneity in *σ* for males, with one particular individual having a very long-distance spatial recapture and many individuals having no spatial recaptures. The samples for this individual were rarely combined into one individual until 3 loci were used and as *n^cap^* = 37 (the correct number of capture males) became increasingly probable with the addition of more loci at which point, the estimate of *σ* converged upwards to the full SCR estimate. This behavior is consistent with the simulations where *σ* is large, but is more pronounced in this data set, which could be explained by individual heterogeneity in the detection function parameters. A second factor that tends to split the samples from this potentially large *σ* individual apart is that there were several traps between the two traps where this individual was captured and the categorical SPIM found it unlikely that this individual would not have been captured at these traps closer to its estimated activity center until enough categorical covariate information was available to make it more even more unlikely that two individuals in the population had the same multilocus genotype. The second longest spatial recapture in the male data set spans a gap with no traps and required fewer loci to reliably link its samples together.

The posterior distributions of *n^cap^* in Figure 6 demonstrate what we believe is a source of the positive bias in the categorical SPIM estimator. With the addition of just 2 loci for females and 1 locus for males, all incorrect combinations of samples were ruled out by the genotype information and the spatial distribution of the samples. However, a 1 or 2 loci genotype is not sufficient to guarantee the local uniqueness of a genotype, leading to a situation in which samples cannot be erroneously combined into fewer individuals than produced them, but they can be erroneously split apart into more individuals than produced them. Thus, in these scenarios, *n^cap^* can never take a value lower than the true value, but rather must always be larger than the true value. We believe the identity exclusions are removing the lower tail of the posterior distribution of *n^cap^* that would be present in the unmarked SCR estimator, introducing positive bias, which can be removed by adding more categorical identity information and reducing the upper tail of the posterior of *n^cap^*. In the simulations, this occurs with the addition of just a few identity categories; however, individual heterogeneity in detection function parameters as argued above, may require more categorical identity information to remove the positive bias in *n^cap^* and thus *N*.

This analysis also demonstrated the use of genotypes that are partial as a consequence of DNA amplification failure, with two caveats. First, there were very few usable partial genotypes because of the DNA amplification protocol used in which samples at the same trap/occasion were subsequently genotyped until a full genotype was obtained. This process led to the partial genotypes matching the complete genotype individual at a particular trap/occasion with high probability because bears usually leave multiple hair samples in a hair snare, violating the Bernoulli observation process. Second, we assumed the partial genotype samples did not contain any genotyping errors. Three of the 6 partial genotype samples used matched other individuals in the population with high probability, but 3 partial genotypes had posterior probabilities of 1 that they were new individuals. These may have indeed been new individuals, or perhaps they did not match any other individuals because they were corrupted. Including partial genotypes in this manner needs to be done with caution and in consultation with a wildlife geneticist, or the categorical SPIM could be extended to accommodate genotyping errors (e.g. Wright *et al*., 2009). If partial genotypes, or even a subsample of the partial genotypes, can be deemed reliable, including them in the analysis can increase the precision of abundance and density estimates, especially if high probability spatial recaptures can be added, as was the case in the female bear data set.

## 5 Discussion

We developed a spatial capture-recapture model for categorically marked populations that uses any number of partially identifying categorical covariates to reduce the uncertainty in the individual identity of latent identity samples via three mechanisms. First, any samples that are inconsistent at any observed covariates are deterministically excluded from matching. Second, as the number of identity categories created by covariates increases and as the category level probabilities for each covariate become more equal, it is increasingly unlikely that more than one individual locally, and in the population, will have the same full categorical identity. Third, the spatial location of the latent identity samples and the estimated detection function scale parameter, *σ*, spatially restrict which samples matching at all observed covariates could have been produced by the same individual. Thus, the categorical SPIM reduces uncertainty in the individual identity of latent identity samples by providing deterministic identity exclusions and reducing the uncertainty in probabilistic identity associations using both spatial and categorical covariate information. The categorical SPIM simulation and MCMC functions are maintained in the SPIM R package (Augustine, 2018) and can also be found in Supplement 2.

A specific case of categorically marked populations that is of some practical importance is that in which individuals are marked by multilocus genotypes. In this case, each locus of a genotype is a single categorical covariate and the categorical SPIM provides a continuous model for genotype uniqueness. Thus, the categorical SPIM is an alternative to using the P(ID) and P(sib) criteria currently used that allows for uncertainty in individual identity as might be the case when fewer loci are amplified than necessary to meet probability of identity criteria, which might occur in populations with very low genetic diversity (e.g. McCarthy *et al*., 2009). The categorical SPIM also introduces the possibility of using fewer loci than necessary to meet probability of identity criteria by design, trading some certainty in individual identity for lower genotyping costs. Genotyping costs do not increase linearly with the number of loci and Puckett (2017) found that variability in the number of loci used in microsatellite studies explains very little variability in total project costs. There may be some cost savings if using fewer loci allows for the use of fewer multiplex panels, which explain a moderate amount of variability in total project costs (Puckett, 2017).

Simulation scenarios A1-4 (Figures 2 and 3) show the importance of population abundance, density given abundance, and *σ* for the accuracy and precision of the unmarked SCR and categorical SPIM estimators. Estimates from populations with lower density given abundance and smaller σs showed less bias and estimates were more precise for populations with higher abundances, lower density given abundance, and smaller σs. More categorical identity groups were necessary to maximize precision when abundance was higher and *σ* larger; however, for the scenario that raised *D* without raising *N*, the majority of precision gains came from the addition of the first 4 identity categories. This scenario raising *D* without raising *N* demonstrates the importance of disentangling the relationship between *N* and *D* when assessing the performance of unmarked SCR, and SCR models with latent or partial individual identities, more generally. Further, it suggests that categorical identity covariates are more effective in populations where estimator uncertainty is due to a large *D* relative to *N*, rather than a large *σ*.

The IDI correlated negatively with the precision and accuracy of *n^cap^* estimates. For the scenarios that increased the IDI holding *N* fixed (A1 vs. A3, A2 vs. A4, A2 vs, A2b), the IDI also correlated negatively with the accuracy and precision of *N* estimates. In scenarios that increased *D* by increasing *N* (A1 vs A2, A3 vs. A4), the increase in uncertainty in *n^cap^* was outweighed by the decrease in data sparsity and *N* estimates were more precise and accurate. Our exploration of IDI values here is very limited and a larger simulation study is necessary to determine how well this index correlates with estimator performance and to what degree do scenarios with differing population density and *σ* values producing the same index value share the same estimator performance. Our results suggest that scenarios with differing *D* and *σ* values that produce the same IDI values will not necessarily produce the same precision in *n^cap^*. We speculate that this is related to how the latent identity samples interact with different densities of activity centers in the SCR process model. Specifically, when *D* is large, there are necessarily more nearby individuals that a latent identity sample can be allocated to, which increases the uncertainty in individual identity. Finally, note that some of the largest values of the IDI may represent ecologically implausible or even impossible scenarios since home range size generally varies inversely with density (Efford *et al*., 2016).

It is not widely recognized that SCR models with latent individual identities where *n^cap^* is latent, specifically, unmarked SCR and SMR, produce biased abundance estimates when data are sparse or that this bias is magnified when density is higher for a fixed abundance and/or *σ* is larger. We demonstrated these effects for unmarked SCR; however, these patterns should also be present in SMR, at least those with very few marked individuals and/or sparse detection data for the marked individuals as can occur when using natural marks and/or surveying low density populations. In fact, even though our simulation specifications were more challenging than those of Chandler & Royle (2013), many studies in practice use fewer than 81 traps that we considered and an expected 1.65 captures per individual (including the individuals captured 0 times) is likely optimistic for some sampling scenarios.

The addition of marked individuals should allow for more reliable estimation for the unmarked population component because the marked individuals provide more information about the detection function parameters and by construction, reduce the number of samples with latent or partial individual identifications. Further, the use of individual-linked telemetry data and a marking process capture history in conjunction with generalized SMR (Whittington *et al*., 2016) improves the estimation of model parameters and thus, the reliability of density estimates. Still, until some practical guidelines can be established, relating estimator performance to abundance, density given abundance, and metrics of data sparsity (e.g., *λ*, *K*, number/spacing/extent of traps), we recommend researchers conduct simulations with unmarked SCR or SMR parameter values appropriate to their study design to determine if their study designs are sufficient to produce reliable estimates using these models. If not, the categorical SPIM offers a second route to remove bias and increase precision via reducing the uncertainty in individual identity, with the first route being the reduction of uncertainty in *σ* using telemetry data (e.g., Sollmann *et al*., 2013) and/or informative priors (Chandler & Royle, 2013; Ramsey *et al*, 2015).

Simulation scenarios B1-3 (Table 1) demonstrate a proof of concept for using data sets where some or most of the full categorical identities are partial such as partial genotypes. With a 0.5 probability of successful amplification at each locus, no data sets produced enough full genotype samples that would have been usable in a model that required certain individual identification, but the categorical SPIM produced an estimate that was 95% as precise as the estimate where all loci were amplified with probability 1. Even assuming only 75% of the samples were usable, the estimate was 80% as precise. Therefore, the categorical SPIM provides a way to use the partial identity information such as partial genotype samples that are currently being discarded. We caution, however, that if partial genotype samples are more likely to have genotyping errors, the categorical SPIM needs to be extended to accommodate those errors, or perhaps a subset of the partial genotype samples could be deemed reliable through consultation with a wildlife geneticist. For example, allelic dropout could be ruled out by discarding any loci that were homozygous or using appropriate tests (e.g., available in MicroChecker, Van Oosterhout *et al*., 2004), or partial genotypes could perhaps be deemed reliable if they repeatedly produced the same partial genotype after a sufficient number of amplification attempts in a multi-tubes approach. Alternatively, adapting the allelic dropout model of Wright *et al*. (2009) to the categorical SPIM would be straightforward and Wang (2017) present a model for false alleles that could be adopted if sufficiently general. Both of these error processes would require the modeling of replicate amplification attempts rather than the consensus genotypes used here.

We will discuss four assumptions of particular importance that we make for the categorical SPIM. First, and perhaps most consequential, is that there is no individual heterogeneity in detection function parameters or similarly, there is no transience in the activity centers during the time of the survey (Royle *et al*., 2016). Like unmarked SCR and spatial mark-resight (SMR), the categorical SPIM uses the detection function likelihood to determine how likely it is that each latent identity sample came from each individual. As demonstrated in Appendix A, of the detection function parameters, *σ* largely determines the degree to which samples from different individuals overlap in space. If there is individual heterogeneity in detection function parameters, especially *σ*, and the SCR model does not include this individual heterogeneity, an average *λ*_0_ and/or *σ* will be estimated, except the average will be biased upwards since the samples will disproportionately belong to the more detectable individuals. Still, the latent identity samples from the most detectable individuals will tend to be incorrectly split across two or more latent individuals because their true spatial distribution of samples will be deemed unlikely by the model based on the averaged *λ*_0_ and/or *σ* values. Therefore, *n^cap^* will tend to be biased high, introducing positive bias into *N*. The black bear example suggests that individual heterogeneity in detection function parameters (if it was present) can be overcome with increasing identity category information to correctly reproduce the true *n^cap^*, but adding individual heterogeneity to the categorical SPIM detection model (as well as unmarked SCR and SMR) would be required to get appropriate abundance estimates. This should be investigated in the future; although, we expect the introduction of individual heterogeneity in detection function parameters to drastically increase the uncertainty in individual identity and thus, the utility of the categorical SPIM, unmarked SCR, and SMR for density estimation. A better strategy would be the use of covariate-specific detection function parameters for specific covariates such as the sex, if the subsets of the population that have different detection function parameters can be at least partially identified.

The second assumption of note is that all possible category levels are known, which may not be the case for genotypes. There are two ways in which genotypes can be identified and enumerated for the categorical SPIM: identifying all observed genotypes at each locus, or identifying all implied genotypes at each locus based on the observed alleles at each locus. The latter method is certainly more thorough, but given sufficiently large data sets, any unobserved genotypes will occur in the population very rarely. Regardless, there will always be the possibility that very low probability genotypes exist in the population that were not observed in the data set, and not accounting for these individuals will introduce negative bias in abundance estimates (Wright *et al*., 2009), though, the magnitude of bias will be small if the majority of genotypes with non-negligible frequencies are identified. Wright *et al*. (2009) raise the possibility of using independent, reference genotypes from the same population to improve abundance estimates. This information could aid in identifying all possible genotypes for each locus and could also be used to aid the estimation of genotype frequencies, which would be especially helpful for sparse data sets. The latter use would require that the reference genotypes were representative of the population subject to capture but did not contain any of the same individuals which would violate independence (Wright *et al*, 2009).

The third assumption of note is that the full categorical identities are independent among individuals. When using genotypes, this assumption will be violated if genetic structure exists at the home range level as a result of relatedness, natural or anthropogenic impediments to movement, or other factors. Due to the importance of the spatial proximity of samples in the categorical SPIM, the combination of relatedness and philopatry (e.g. female black bears) could lead to a spatial correlation in genotypes across the landscape. This could introduce negative bias if sufficiently strong because latent identity samples of nearby, related individuals will be erroneously combined into one individual too often when using few loci data sets. We suspect spatial correlation in genotypes at the home range level will be weak in most spatially structured, non-isolated populations, and the categorical SPIM to be robust to this effect; although, this might not be the case for species such as canids that travel in packs that are highly related. As the spatial uniformity of activity centers can be regarded as a weak prior for the distributions of individuals across the landscape (Royle *et al*., 2013), the spatial uniformity of category levels across the landscape is likely a similarly weak prior with the posterior able to deviate substantially from spatial uniformity, given sufficient data. Regardless, the positive bias from individual heterogeneity in detection function parameters will likely outweigh any negative bias from the non-independence of genotypes, but this effect should be further investigated. The robustness of the categorical SPIM to assumption violations in typical genetic data sets can be further established by seeing how well it can reproduce estimates from full data sets as we did in the black bear application across many species and study areas.

The fourth assumption we will discuss is that the categorical covariates are independent of one another. When using genotypes, this is equivalent to linkage equilibrium, which can be tested for (Rousset & Raymond, 1995), and a test for linkage disequilibrium in the black bear data set used in our application failed to detect linkage disequilibrium (Murphy *et al*., 2016). Non-independence of the categorical covariates may occur for other types of data; for example, when using natural marks, body size and sex may not be independent. In this case, a composite covariate that combined body size and sex could be constructed, but this would require the categorical SPIM to be modified to handle missing data if sometimes only body size or sex was observed. When present, linkage disequilibrium causes pseudoreplication in genetic data sets (Selkoe & Toonen, 2006), which we expect to introduce negative bias into the categorical SPIM estimator because there will be less variability than expected in the updated latent full categorical identities across MCMC iterations.

We added categorical covariates to the unmarked SCR model, but categorical identity covariates can also be added to SMR, 2-flank SPIM, and likely other types of specialized SPIMs that may not have yet been developed. Further, the categorical SPIM as described allows continuous covariates if they are discretized, which requires the assumption that they are measured without error. Continuous covariates with measurement error models could be added for a more general “covariate SPIM”. In fact, the spatial location of a sample is a continuous covariate already being used by SPIMs, and the SCR process and detection function models can be conceptualized as a measurement error model for the individuals’ activity centers where the variance of the measurement error process is largely determined by the size of an animal’s home range. An alternative, but likely less efficient way to accommodate measurement error is to classify borderline observations as missing data. For example, if one can ascertain body size from remote cameras, they may be classified as small and large, with intermediate observations classified as missing data. Another ecologically relevant continuous covariate that is informative of identity is the time a sample was recorded with samples collected closer together in time being more likely to have been produced by the same individual. However; making use of the time of sample deposition would require a model for animal movement and is probably less informative than spatial location, unless *σ* is large.

While our application and much of the discussion focuses on the use of microsatellite loci as categorical identity covariates, other observation systems provide categorical identity covariates that can be used by the categorical SPIM. Remote cameras, for example, sometimes provide individual sex, age class, color morph, natural markings, and/or other morphological features (Villafahe-Trujillo *et al*., 2018). Some species such as mustelids (Royle *et al*., 2011; Villafane-Trujillo *et al*., 2018; Siren *et al*., 2016) and Andean bears (Molina *et al*., 2017) can have markings on the chest, head, and neck, that can be used to classify unique individuals. We suspect categorical and/or continuous covariates can be extracted from these types of observations that would allow this identifying information to be used without having to assign a unique identity with certainty, which could be erroneous if two individuals shared the same markings, or to include the photographs that were not distinct enough to provide a unique identity. Further, if hair snares and remote cameras are deployed together (e.g., Royle *et al*., 2011) categorical identity covariates from photographs can be combined with those provided by microsatellite loci.

In addition to the possible ways we envision the categorical SPIM may be used outlined in this paper, it may allow for better density estimation for currently used observation systems we are not aware of, and may spur the adoption of new ways of categorically marking species for which unique marks are not currently available once researchers are aware that density can be estimated using the information contained in categorical marks. Unique marks will always be the most informative, but the categorical SPIM offers a middle ground between uniquely marked and unmarked populations. SMR is another SCR-based model that in intermediate between fully known and full latent individual identities and combining the categorical SPIM with SMR is especially appealing because it would allow all of the features currently available to improve density estimates over unmarked SCR to be combined into a single model that can accommodate a subset of marked individuals, individual-linked telemetry data, a marking process capture history, and categorical identity covariates.

## Tables and Figures

**Table 1:**
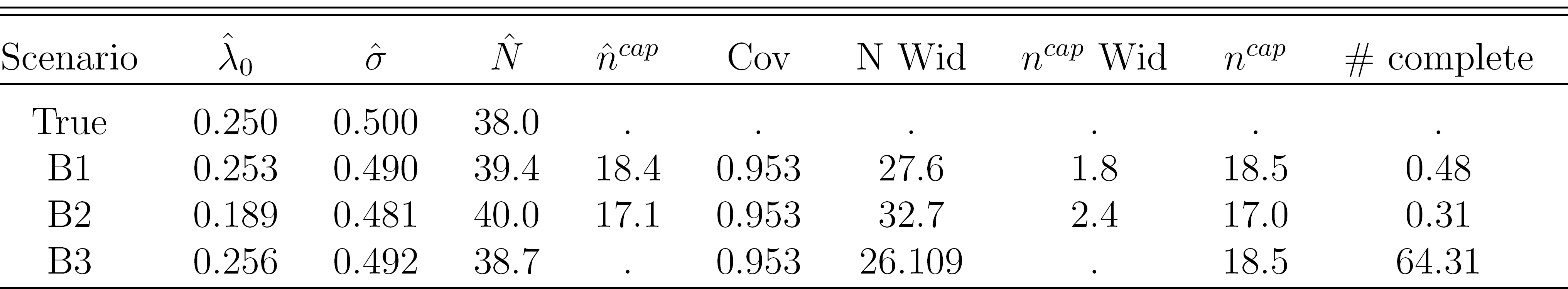
Simulation results for the partial genotype analyses. Scenarios B1 and B2 had loci amplification probabilities of 0.5. Scenario B1 used all partial genotype samples, while Scenario B2 used 75% of the partial genotype samples. Scenario B3 is a typical SCR model using all full genotype samples for comparison. Parameter mean point estimates are presented, along with coverage of *N* (Cov), the mean 95% credible interval widths for *N* (N Wid) and *n^cap^* (n wid), and the mean number of samples with complete genotypes (# complete).

**Table 2:**
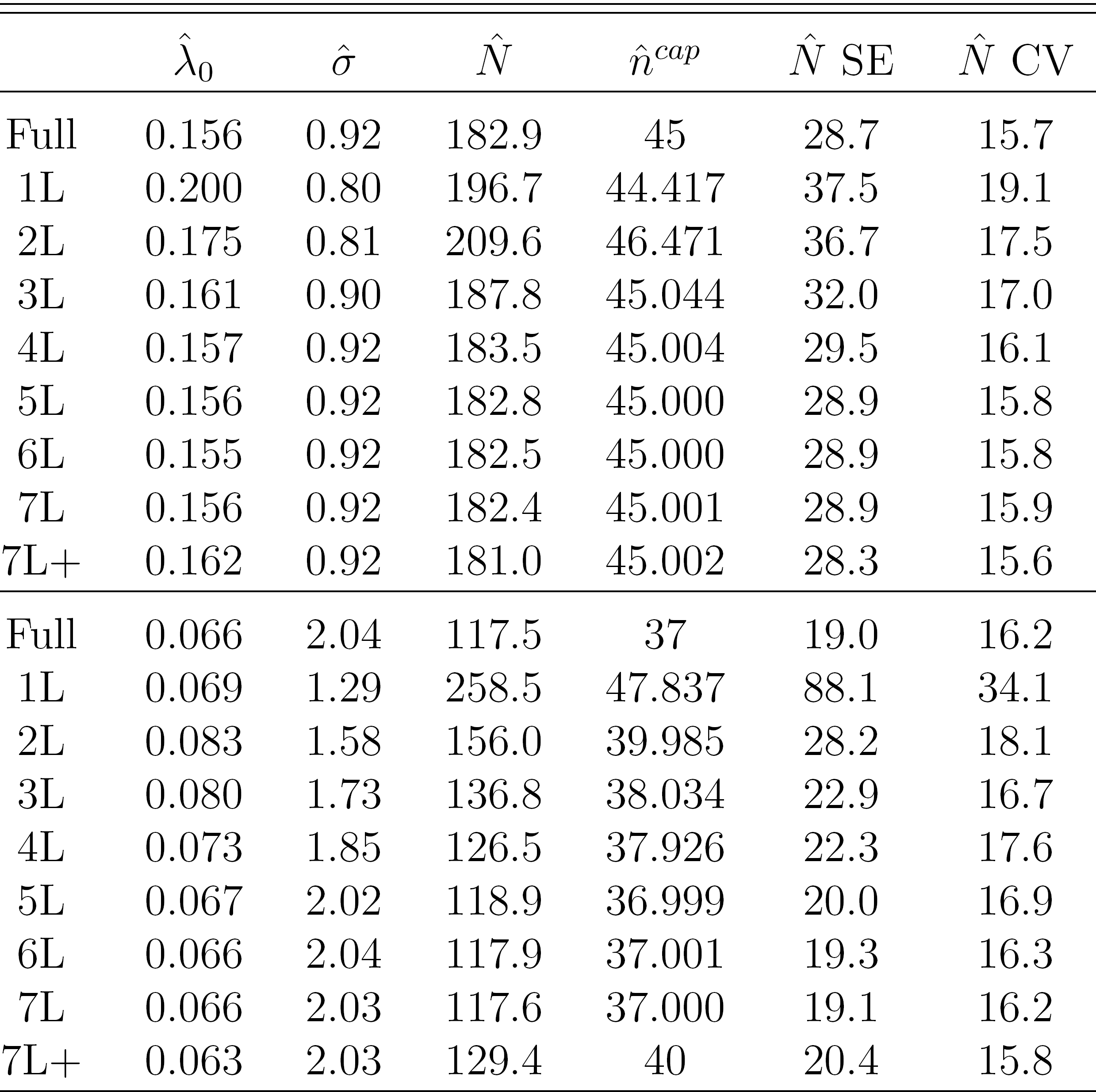
Sex-specific estimates of the detection function baseline encounter rate, λ_0_, the detection function spatial scale parameter, *σ*, and abundance from regular SCR (Pull), categorical SPIM using 1-7 loci (1L – 7L), and categorical SPIM with 7 loci, adding partial identity samples not included in the SCR estimate (7L+). Point estimates are the posterior mode, uncertainty is quantified by the posterior standard deviation (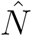 SE) and the coefficient of variation (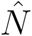 CV). For the categorical SPIM models, 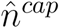 is the estimated number of captured individuals. Female estimates are listed first, followed by males.

| | $\hat{\lambda}_0$ | $\hat{\sigma}$ | $\hat{N}$ | $\hat{n}^{cap}$ | $\hat{N}$ SE | $\hat{N}$ CV |
| --- | --- | --- | --- | --- | --- | --- |
| Full | 0.156 | 0.92 | 182.9 | 45 | 28.7 | 15.7 |
| 1L | 0.200 | 0.80 | 196.7 | 44.417 | 37.5 | 19.1 |
| 2L | 0.175 | 0.81 | 209.6 | 46.471 | 36.7 | 17.5 |
| 3L | 0.161 | 0.90 | 187.8 | 45.044 | 32.0 | 17.0 |
| 4L | 0.157 | 0.92 | 183.5 | 45.004 | 29.5 | 16.1 |
| 5L | 0.156 | 0.92 | 182.8 | 45.000 | 28.9 | 15.8 |
| 6L | 0.155 | 0.92 | 182.5 | 45.000 | 28.9 | 15.8 |
| 7L | 0.156 | 0.92 | 182.4 | 45.001 | 28.9 | 15.9 |
| 7L+ | 0.162 | 0.92 | 181.0 | 45.002 | 28.3 | 15.6 |
| Full | 0.066 | 2.04 | 117.5 | 37 | 19.0 | 16.2 |
| 1L | 0.069 | 1.29 | 258.5 | 47.837 | 88.1 | 34.1 |
| 2L | 0.083 | 1.58 | 156.0 | 39.985 | 28.2 | 18.1 |
| 3L | 0.080 | 1.73 | 136.8 | 38.034 | 22.9 | 16.7 |
| 4L | 0.073 | 1.85 | 126.5 | 37.926 | 22.3 | 17.6 |
| 5L | 0.067 | 2.02 | 118.9 | 36.999 | 20.0 | 16.9 |
| 6L | 0.066 | 2.04 | 117.9 | 37.001 | 19.3 | 16.3 |
| 7L | 0.066 | 2.03 | 117.6 | 37.000 | 19.1 | 16.2 |
| 7L+ | 0.063 | 2.03 | 129.4 | 40 | 20.4 | 15.8 |

**Figure 1:**
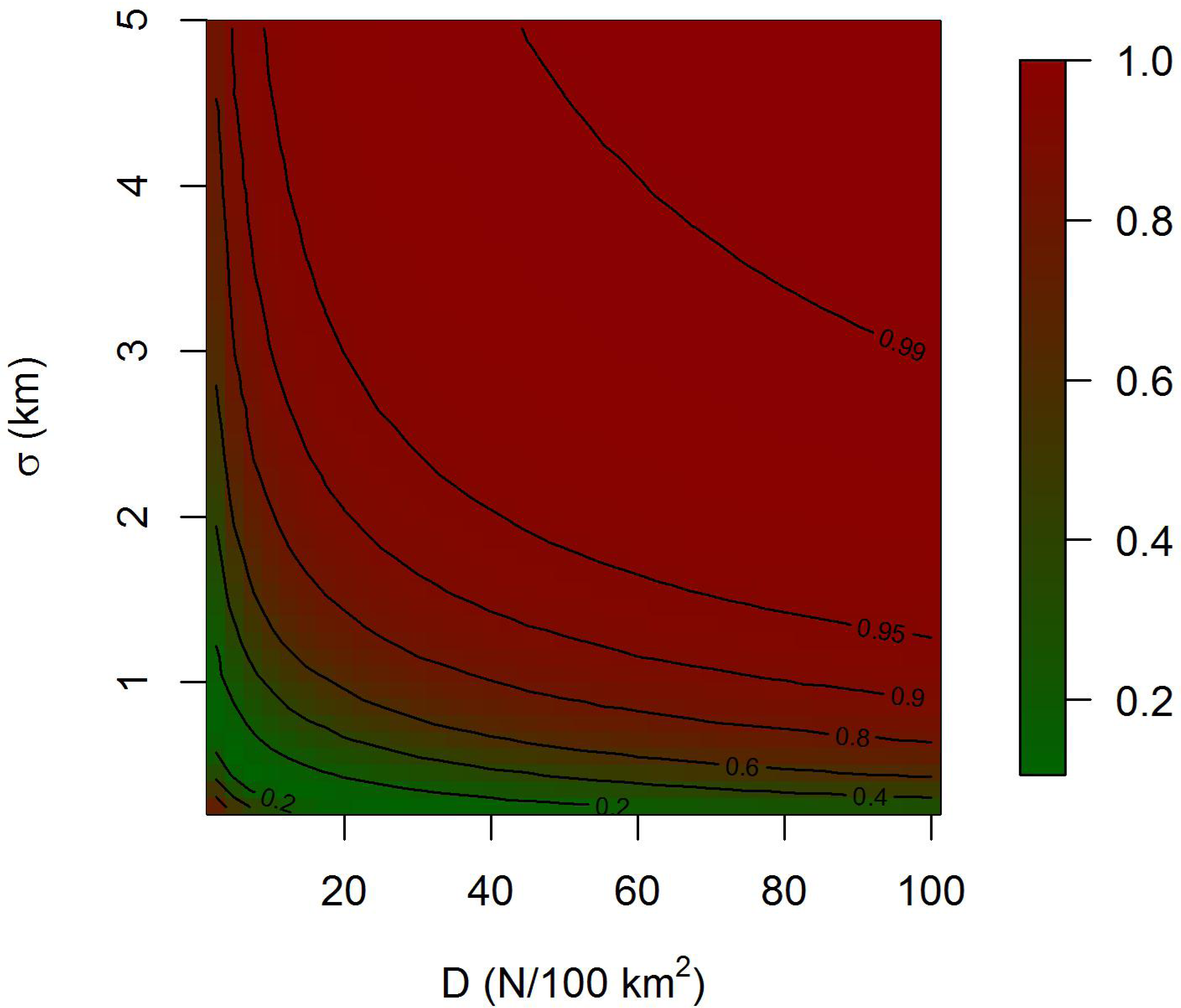
The Individual Identity Diversity Index, quantifying the magnitude of uncertainty in individual identity, as a function of population density (D) and the SCR spatial scale parameter (σ). Increasing D and *σ* increases the magnitude of uncertainty in individual identity and with *N* fixed, the expected precision of *N* and *D* estimates.

**Figure 2:**
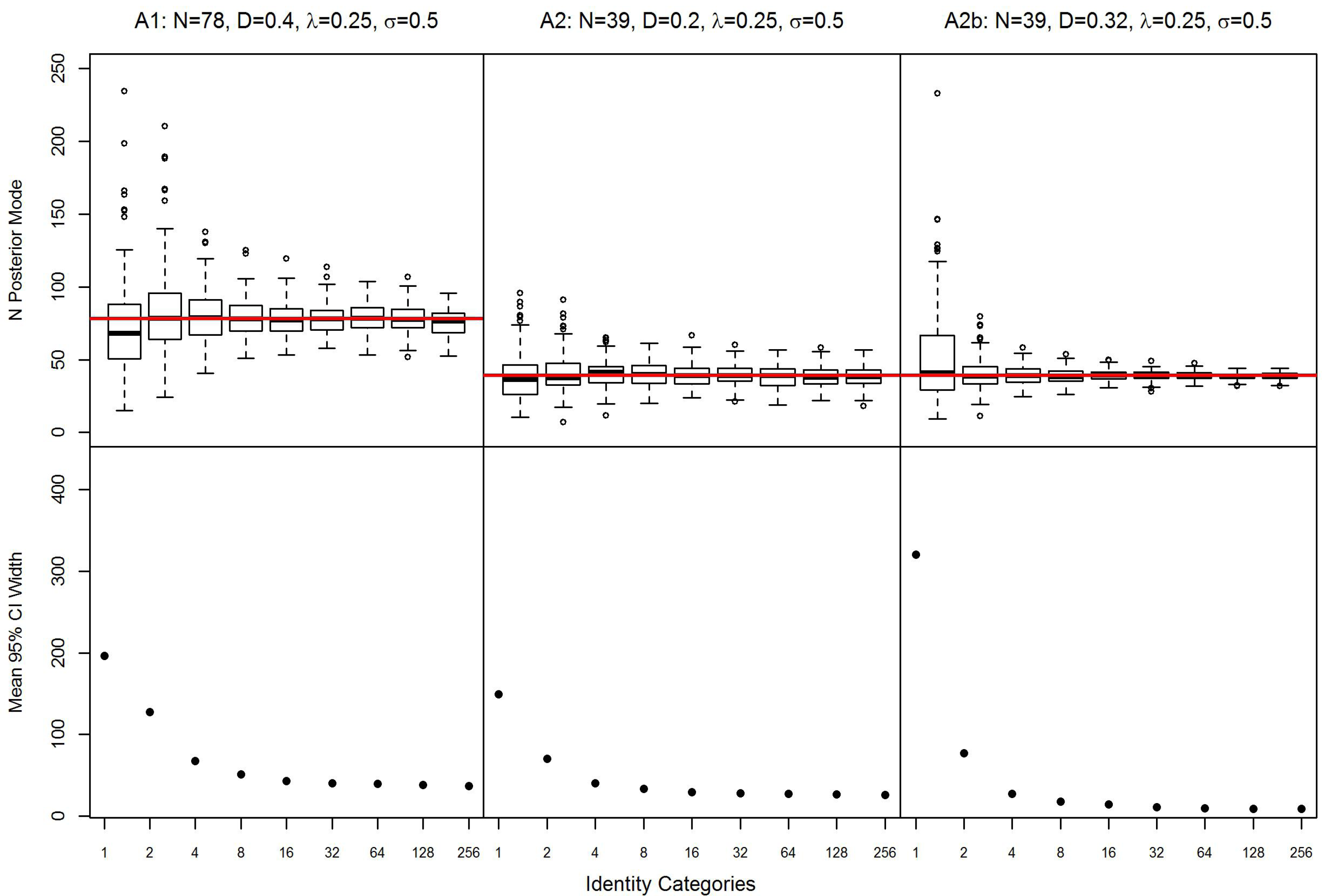
Posterior modes and mean 95% credible interval width of abundance (N) estimates plotted against the number of identity categories in Scenarios A1, A2, and A2b with baseline detection rate *λ*_0_ = 0:25 and detection function spatial scale parameter *σ*=0.5. Population Density, *D* varies across scenarios, and A2b is the scenario that increases *D* while *N* remains fixed. Note the number of identity categories increase exponentially.

**Figure 3:**
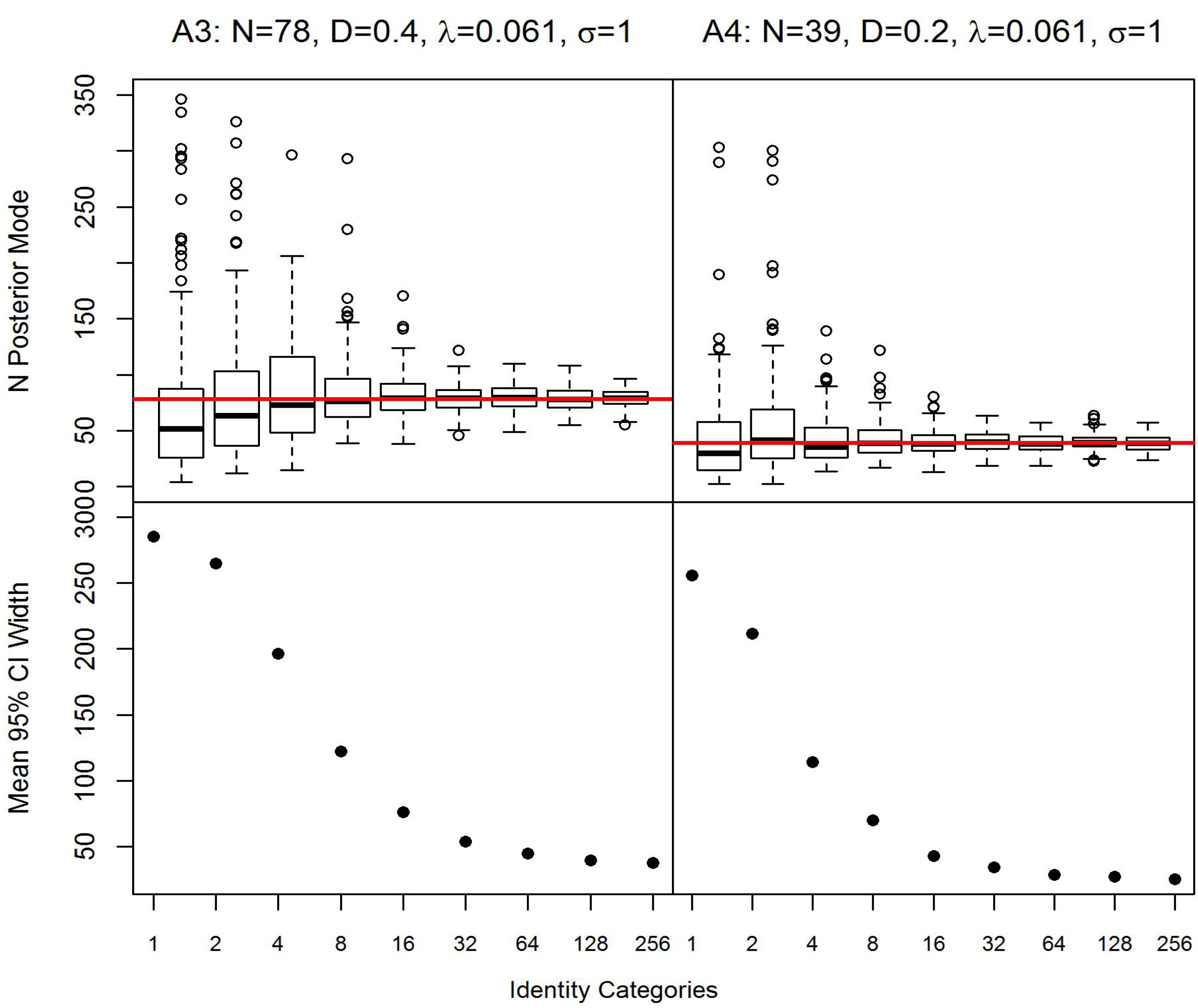
Posterior modes and mean 95% credible interval width of abundance (N) estimates plotted against the number of identity categories in Scenarios A3, and A4 with baseline detection rate *λ*_0_ = 0.061 and detection function spatial scale parameter *σ*=1. Population Density, *D* varies across scenarios. Note the number of identity categories increase exponentially.

**Figure 4:**
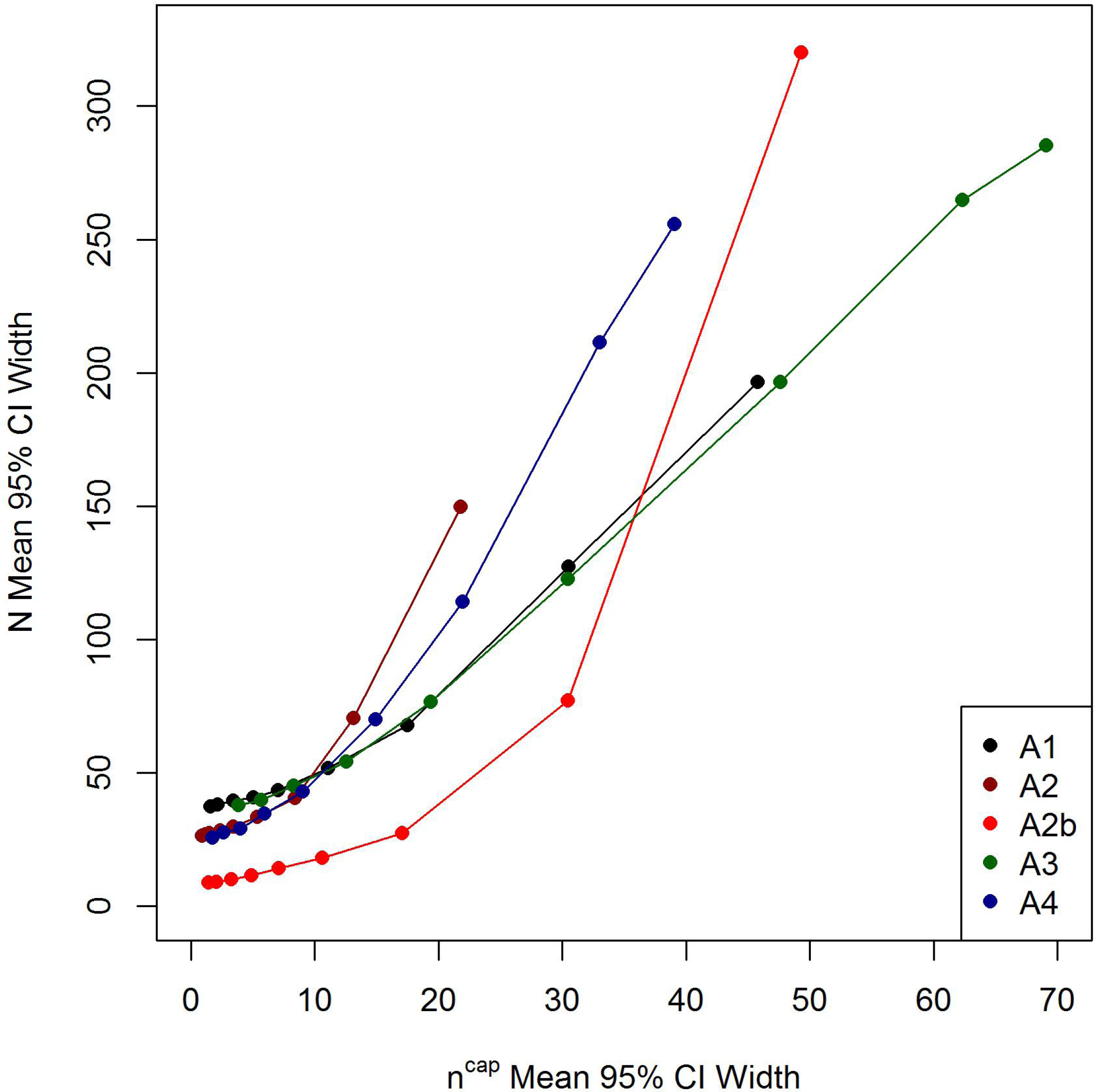
The relationship between uncertainty in the number of individuals captured, *n^cap^* and uncertainty in abundance, *N* for each of the 5 A scenarios.

**Figure 5:**
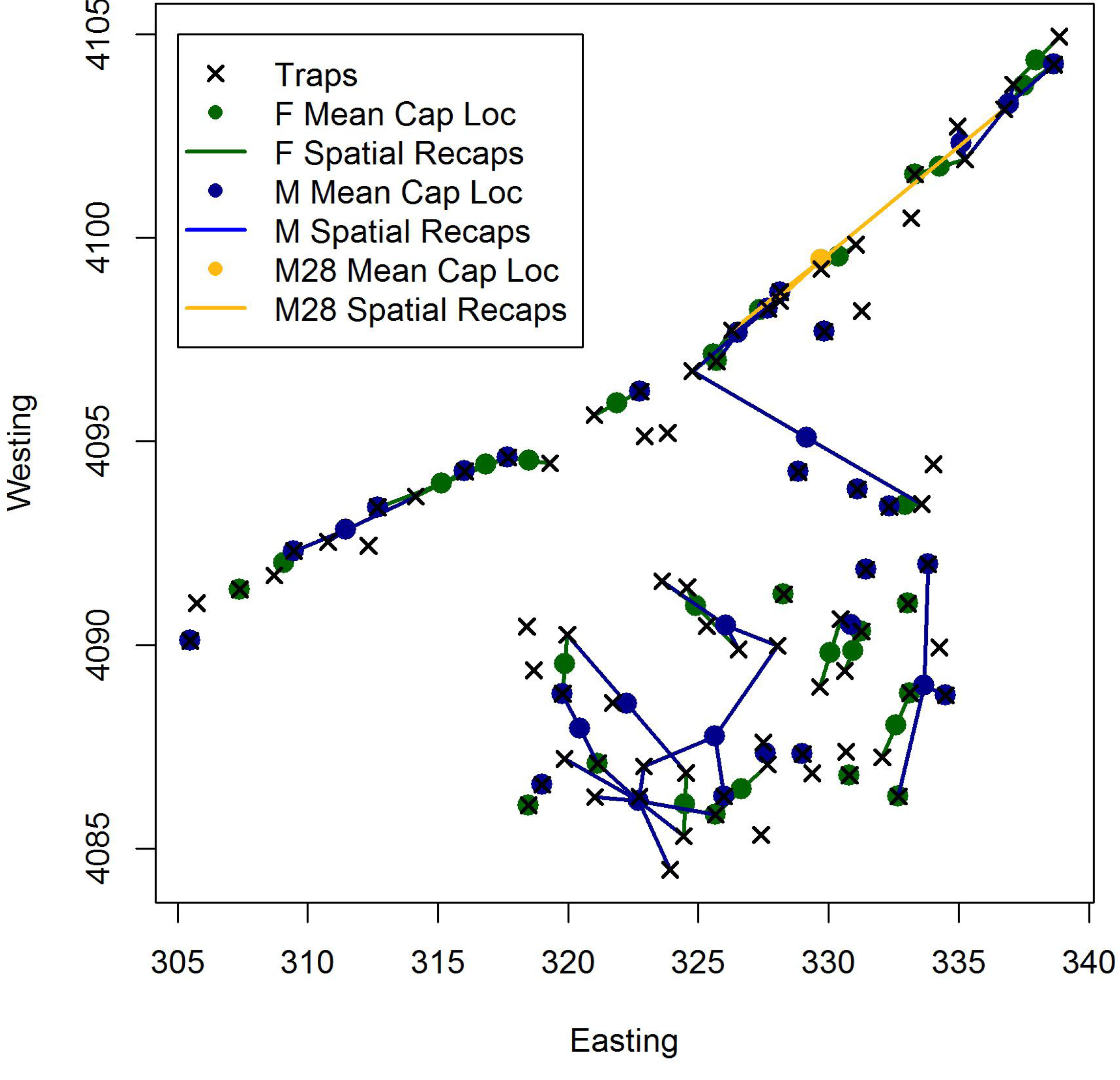
Mean male and female capture locations with spatial recaptures for the Kentucky black bear data set. Male 28 is highlighted in yellow because it had a large spatial recapture and was not detected at several traps in between the two traps where it was detected, requiring more loci for the categorical SPIM to link these two samples together.

**Figure 6:**
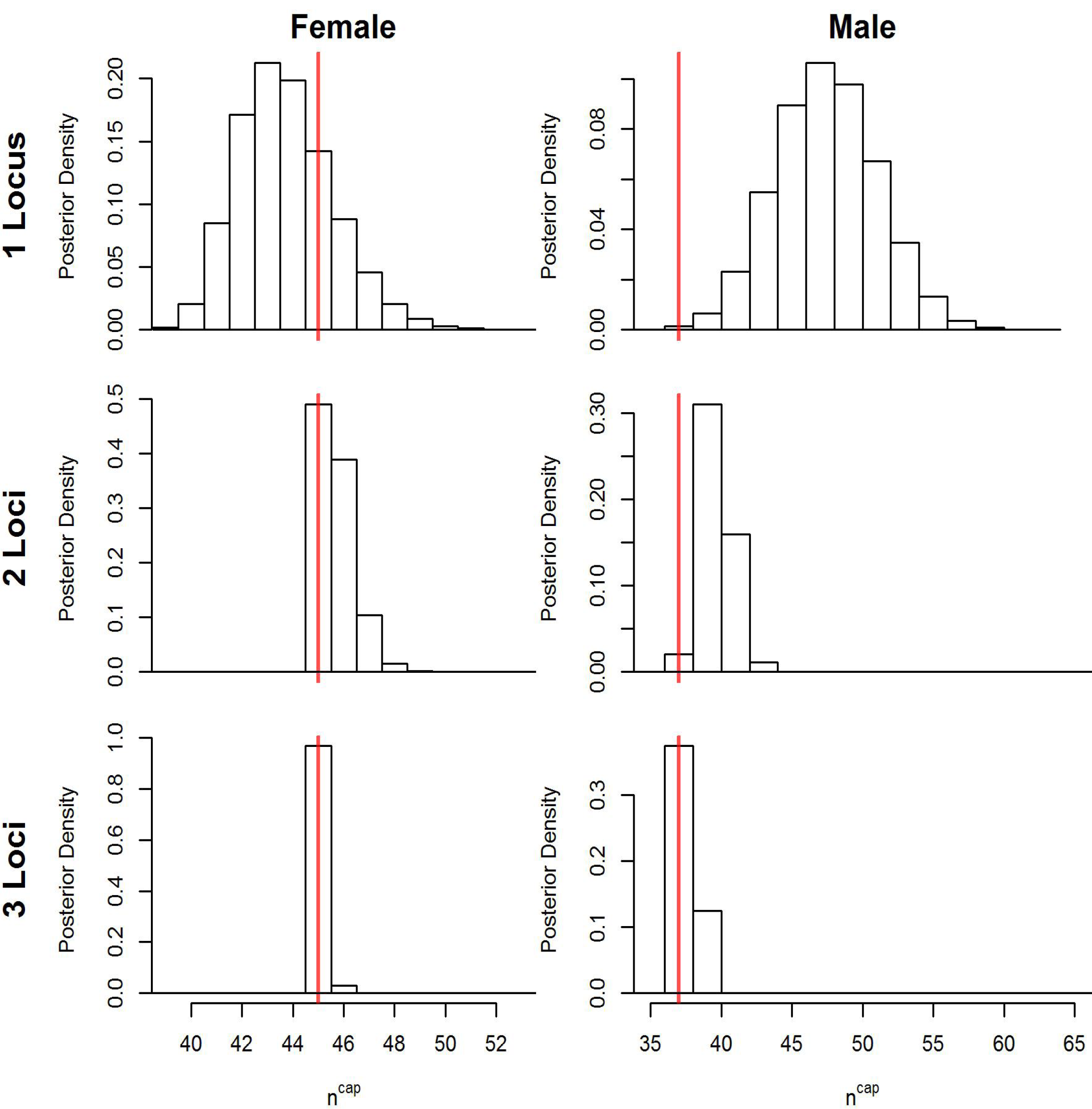
Posterior distributions for the number of unique individuals captured (*n^cap^*) for the Kentucky black bear female and male data set using 1, 2 and 3 loci. True values from full data sets are marked in red (45F, 37M).

## 7 APPENDIX A: Individual Identity Diversity Index

We propose using the Simpson Diversity Index (Simpson, 1949) to quantify the magnitude of uncertainty about individual identity in models with latent or partially latent individual identities. The general idea is that there will be more uncertainty in individual identity of the latent identity samples when there are more individuals exposed to capture and when more individuals have closer to an equal probability of being captured at any point on the landscape if a trap were to be placed there. Population abundance determines how many individuals are exposed to capture at each point on the landscape-all *N* individuals are exposed to capture at all points on the landscape, even if the detection probability of most individuals is effectively zero. Population density, *D*, and the detection function spatial scale parameter, *σ*, then determine how many individuals will have non-negligible capture probabilities and/or capture probabilities of similar magnitudes at any point on the landscape. The Simpson’s Diversity Index can be applied to quantify this diversity of individual identity at any point on the landscape as a metric of the magnitude of uncertainty in individual identity of the latent-identity samples that is independent of the observation process that includes the trapping array and number of trapping occasions and, as we will show, is negligibly affected by the value of the detection function baseline detection parameter, *λ*_0_.

The Identity Diversity Index (IDI) can be calculated at each trap conditional on one realization of the activity centers; however, a single metric for each combination of *D*, *λ*_0_, and *σ* can be achieved using the expected ID averaged over many traps and realizations of the activity centers. The expected ID can be calculated at any point on the landscape, whether or not a trap is located there by discretizing the state space into *J* grid cells of equal size. We define the individual by grid cell level detection probability of the binomial observation model to be *p_ij_, i* = 1, *N*, with *p_ij_* determined by the distance between activity center *i* and spatial location *j* and a detection function, such as the hazard half-normal described in the Methods. We then normalize these probabilities so they sum to 1 at each trap by 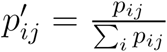. Then ID at spatial location *j* is 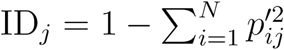 which is exactly one minus the Simpson’s Diversity Index so that larger values indicate more uncertainty in individual identity. If the spatial distribution of activity centers across the landscape is uniform, the expected value of ID will be the same in all grid cells, except there is an edge effect induced by the edge of the state space, which we removed by buffering the grid area by at least 3 times the detection function scale parameter, *σ*. For each combination of *D*, *λ*_0_, and *σ* (ranges listed below), we simulated K=25 realizations of the activity centers on a state space of size 1444 km^2^ (a 20 × 20 km grid buffered by 3 times the maximum *σ)* and calculated *ID_jk_* on the inner 400 km^2^ divided into 1681 grid cells. The expected ID was then estimated by averaging *ID_jk_* over the 25 activity center replications and 1681 grid cells. For simplicity in the main text and in plots, we refer to the expected ID as the Identity Diversity Index.

To first demonstrate that *λ*_0_ has a relatively minor effect on expected ID, we calculated the Identity Diversity Index for all combinations of *D* ranging from 5 to 50 N/100 km^2^ in increments of 0.05, *λ*_0_ ranging from 0.025 to 0.25 in increments of 0.025, and *σ* ranging from 25 to 3 km in increments of 0.25. We found no evidence of an interaction between any of these parameters, so Figure A1 shows the relationship between each parameter and the Identity Diversity Index plotted at the median values of the other parameters. Note that the Identity Diversity Index does not vary noticeably across the range of *λ*_0_ we considered here,which covers much of the range seen in practice, although there is a very slight increase of size 0. 01 in the Identity Diversity Index as *λ*_0_ is increased across the full range of *λ*_0_ considered here. This minimal change is because the distribution of *p_ij_* at location *j* is shifted right when *λ*_0_ is raised and normalizing produces roughly the same values of 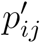 for different values of *p_ij_*.

We then calculated the Identity Diversity Index across a wider range of parameter values for *D* and *σ* at more granular level. For this second set of simulations, *D* ranged from 2.5 to 100 N/100 km^2^ in increments of 2.5, *σ* ranging from 0.25 to 5 km in increments of 0.1 and λ was set to 0.025. We used the same 20 x 20 inner grid to calculate the Identity Diversity Index, but the state space area was larger, at 2500 km^2^, due to the larger maximum sigma. These simulations are presented in Figure 1 of the main text.

**Figure A1:**
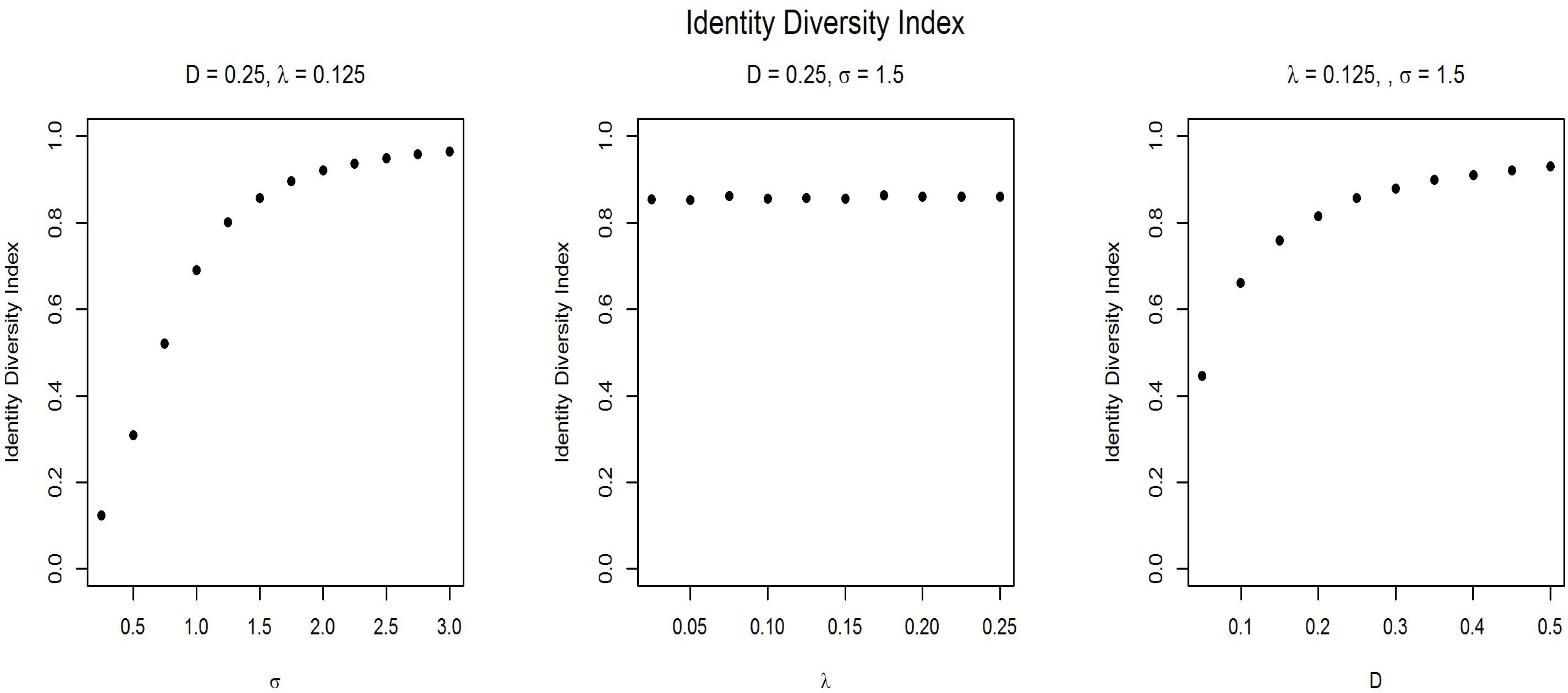
The relationship between the Identity Diversity Index and population density, *D*, and the spatial capture-recapture baseline detection parameter, *λ*_0_, and scale paramater, *σ*.

## 8 APPENDIX B: MCMC algorithm

The inferential goal is to probabilistically reconstruct the true capture histories and each individual’s full categorical identity from the observed trap-referenced detections and associated categorical identities, with potentially missing values, and then produce estimates of abundance and density that incorporate the uncertainty stemming from imperfect individual identity. Here, we will restate some of the notation outlined in the Methods. The observed trap-reference detections are stored in the *n^obs^* × *J* matrix ***Y***^obs^, a matrix of all zeros, except for 1 elements indicating which trap each of the *n^obs^* detections were observed at. The observed categorical identities are stored in the *n^obs^* × *n^cat^* matrix ***G**^obs^*. The true capture history and associated full categorical identities are then stored in *Y^true^* and ***G**^true^*, respectively, both with *M* rows after data augmentation and the same number of columns as their observed counterparts. The process model quantities of interest are *γ*, the population frequencies of the identity category levels for each 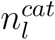 category level for the *l* covariates, ***S***, the individual activity centers, *z*, the data augmentation indicator vector, and *ψ*, the binomial inclusion probability for the data augmentation indicator vector.

The joint posterior is:

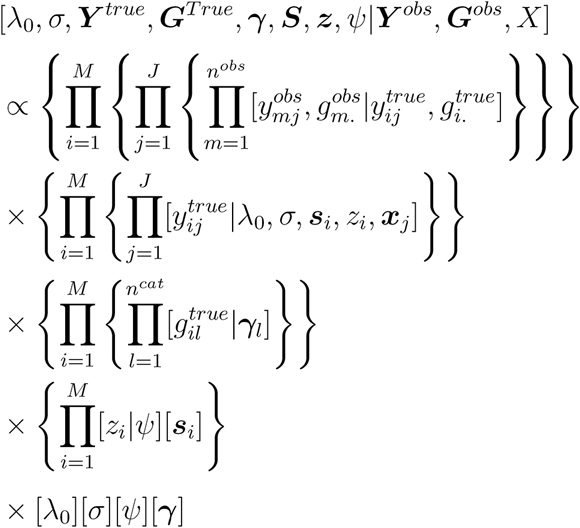

The prior distributions are

1. π(*λ*_0_) ~ Uniform(0, oo)
2. π(*σ*) ~ Uniformed, oo)
3. π*(ψ)* ~ Uniform(0,1)
4. π*r*(*s_i_*) ~ Uniform(*S*)
5. π(*γ_l_*) ~ Dirichlet(*α_l_*), where *oci* is the vector of Dirichlet parameters of length 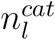, indexed by *g* below. All *α_l_* were set to vectors of 1. Then the joint posterior is the product of the unmarked SCR likelihood, the categorical identity likelihood, and the prior distributions.

Here, we list the full conditional distributions or the distributions that the full conditionals are proportional to. Note the first five distributions are exactly the same as a typical SCR model, the sixth and seventh correspond to the categorical identity component, and the final component corresponds to the uncertain identity component.

1. 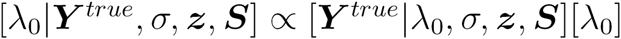, where 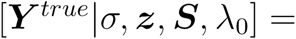 = 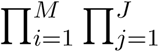 Binomial 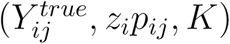 for the Bernoulli observation model and 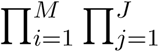 Poisson 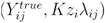 for the Poisson observation model.
2. 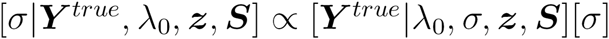 where 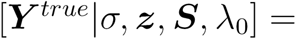 = 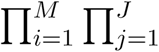 Binomial 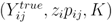 for the Bernoulli observation model and 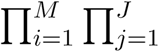 Poisson 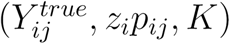 for the Poisson observation model.
3. 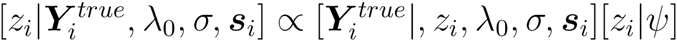, where 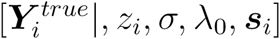 = Bern 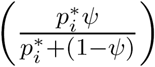 (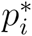 defined below)
4. 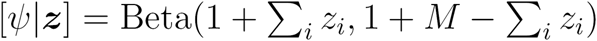
5. 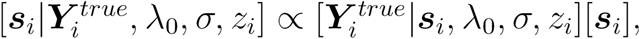, where 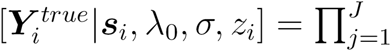 Binomial 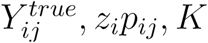 for the Bernoulli observation model and 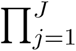 Poisson (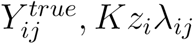) for the Poisson observation model.
6. 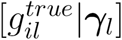 = Categorical (*γ_l_*)
7. 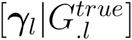=Dirichlet(*y_l_* + *α_l_* where *yl_g_* is the number of individuals in the population (*z_i_* = 1) with categorical identity *g* at loci *l* and *α_l_* is the Dirichlet prior for *γ_l_*, a vector of 1’s.
8. 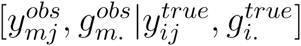. See below.

Here we outline the MCMC algorithm. Again, note that the first five steps are exactly the same as a typical SCR model.

1. Update *λ*_0_. *λ*_0_ is updated with a Metropolis-Hastings step using the distribution Normal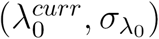, to propose 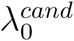, automatically rejecting if a negative value is proposed. The MH ratio is calculated using distribution 1 above.
2. Update *σ*. *σ* is updated with a Metropolis-Hastings step using the distribution Normal(*σ_curr_*, *σ_σ_*), to propose *σ^cand^*, automatically rejecting if a negative value is proposed. The MH ratio is calculated using distribution 2 above.
3. Update **z**. Each *z_i_* is updated by a Gibbs step using the full conditional above where 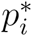 is the probability individual *i* was not captured during the experiment. Let 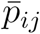 be the probability of not being captured for each individual at each trap on 1 occasion. For the bernoulli observation process, this is 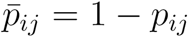 and for the Poisson observation process, this is 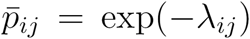. The probability of not being captured during the experiment for each individual is then 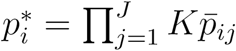.
4. Update *ψ. ψ* is updated with a Gibbs step. Since *π(ψ)* ~ Uniform(0,1) is in the Beta family, the full conditional distribution for *ψ* is Beta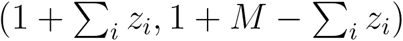.
5. Update **s.** Each activity center *s_i_* is updated with a Metropolis-Hastings step using the distributions Normal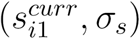 and Normal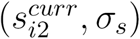 to propose 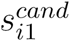 and 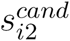, respectively. Proposals that fall outside of the state space are rejected. The full conditional distribution is the SCR observation model likelihood, whether bernoulli or Poisson.
6. Update *Y^true^*. To update *Y^true^* for the Poisson observation model, we use a class-structured version of the algorithm introduced by Chandler & Royle (2013). For the case where all samples are compatible with each other, Chandler & Royle (2013), update the true, latent data at each trap, one at a time, using the multinomial full conditional that follows from contingency table results for Poisson random variables. The full conditional from Chandler & Royle (2013), removing the *k* dimension of occasion, is:

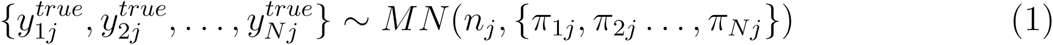

where *n_j_* is the number of samples recorded at trap *j* over all *K* occasions and *π_ij_* = 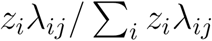. However, if some samples cannot be combined with others as is the case when they have different full categorical identities, the latent true identities of each sample must be updated one by one. Further, the observed data must be organized in a manner that they can be associated with the partially-identifying information such as a genotype. Chandler & Royle (2013) organize the latent identity samples in a vector of trap counts, *n_j_*, or a matrix of trap by occasion counts *n_jk_*, while we organize the latent identity samples with no grouping in ***Y**^obs^*, an *n^obs^* × *J* matrix, which is associated with the observed genotypes, *G^obs^*, an *n^obs^* × *n^cat^* matrix, by the shared *m* dimension spanning 1 … *n^obs^*. We then swap the latent individual identity of each sample one at a time. For each focal sample, *f*, stored in the trap-referenced observed detection vector 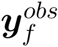 we first identify all individuals currently in the population (*z_i_* = 1) whose latent full categorical identity match the observed categorical identity of the focal sample at all observed covariates (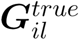 matches 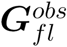 for all *l* that satisfy 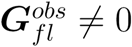, with 0 indicating a missing value), creating an indicator vector for consistency of the full categorical identity, *q*, where *q_i_* = 1 if the latent full categorical identity of individual *i* is compatible with the focal sample’s observed categorical identity and *q_i_* = 0 otherwise. We then choose a new identity for the focal sample with a categorical draw from the probability vector 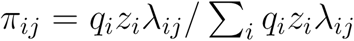, which is identical to how one would update the latent identities in the Chandler & Royle (2013) model, except the probabilities that would swap the latent sample identity to an individual with an inconsistent categorical identity have been zeroed out. Finally, the latent, true data *Y^true^* is updated by moving the focal sample trap-referenced observed detection vector, 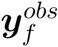, from the current to the new identity (row) of *Y^true^*. The multinomial full conditional result does not hold for the Bernoulli observation model; however, so we use a Metropolis-Hastings proposal. We use the categorical full conditional above as the proposal distribution *π(ID)* ~ Categorical({*π*_1*j*_, *π*_2*j*_ …, *π_Nj_*}, with 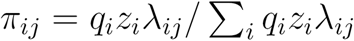. Because the transition distribution for selecting samples to update is not symmetric, the differing forward and backwards transition probabilities must be calculated. The forward transition probability is the probability of selecting a sample to update and then the probability of choosing the new identity *ID*’, *p_for_* = 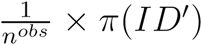. The backwards transition probability accounting for the fact that you can only select individuals with full categorical identities compatible with the focal observed categorical identity to move back is 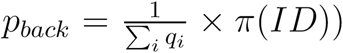. We then accept the proposal with probability:

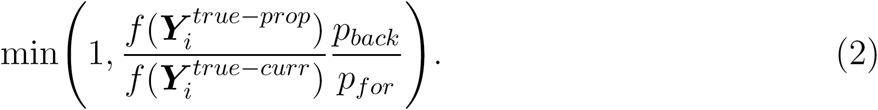

where *f*(.) is the SCR observation model likelihood.
7. Update ***G**^true^*. First, let *G^latent^* be an *M×n^cat^* indicator matrix with elements 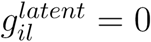 if the observed categorical identity currently associated with individual *i* determine 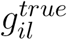 and 1 if not, e.g. all category *l* entries of the samples allocated to individual *i* have an observed genotype of 0 (missing), and are thus consistent with any 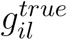 value. Then, *i* and *l* entries of ***G**^true^* are updated with a Gibbs step using the full conditional Categorical(*γ_l_*) if 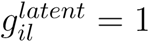 and not updated otherwise.
8. Update *γ_l_*. *γ_l_* is updated with a Gibbs step. Following Wright et al. (2009), we adopt a Dirichlet prior for *γ_l_*, leading to a Dirichlet full conditional with parameter vector 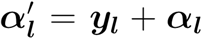 where *y_lg_* is the number of individuals in the population with category level *g* for covariate *l*. To draw values from the full conditional, we first simulate a vector of Gamma random variables *g_l_* ~ Gamma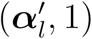, where *g_l_* is of length 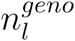. Then, renormalizing these gamma random variables is a draw from the Dirichlet full conditional, i.e. 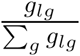.
9. Calculate the derived quantities population abundance, 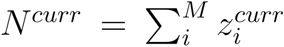, population density, 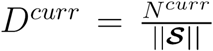, and the number of unique individuals captured *n^curr^* = 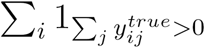

## 9 APPENDIX B: Simulation A Results

**Table B1:** Results from Simulation A Scenarios 1-4. The columns are “IDCat”, the number of identity categories, the mean point estimates, the root means squared error of the point estimates, the coefficients of variation for the point estimates (100xposterior sd/posterior mode), the mean width of the 95% CIs for *N*, the 95% CI coverage of *N*, and *n^cap^*, and the mean number of simulated individuals that were captured. The Identity Diversity Indices for scenarios A1-4 were 0.38, 0.23, 0.76, 0.58. The Individual Diversity Index for scenario A2b was 0.37.

| Scen | IDCat | $\hat{\lambda}_0$ | $\hat{\sigma}$ | $\hat{N}$ | $\hat{n}^{cap}$ | $\hat{N}$<br>RMSE | $\hat{n}^{cap}$<br>RMSE | $N$<br>CV | $\hat{n}^{cap}$<br>CV | $N$<br>Wid | $\hat{n}^{cap}$<br>Wid | $N$<br>Cov | $n^{cap}$ |
| --- | --- | --- | --- | --- | --- | --- | --- | --- | --- | --- | --- | --- | --- |
| A1 | 1 | 0.246 | 0.487 | 75.2 | 37.2 | 40.4 | 13.7 | 75.7 | 34.2 | 196.5 | 45.8 | 0.951 | 36.5 |
| A1 | 2 | 0.243 | 0.482 | 83.0 | 39.0 | 30.8 | 10.6 | 43.1 | 21.1 | 127.4 | 30.5 | 0.917 | 36.9 |
| A1 | 4 | 0.246 | 0.493 | 79.7 | 37.4 | 17.9 | 4.9 | 22.8 | 12.6 | 67.8 | 17.5 | 0.938 | 37.1 |
| A1 | 8 | 0.247 | 0.495 | 79.1 | 37.0 | 13.3 | 3.0 | 17.3 | 8.2 | 51.6 | 11.1 | 0.972 | 37.0 |
| A1 | 16 | 0.249 | 0.494 | 77.5 | 36.1 | 12.4 | 2.1 | 14.8 | 5.5 | 43.5 | 7 | 0.931 | 36.2 |
| A1 | 32 | 0.255 | 0.495 | 78.3 | 36.8 | 10.3 | 1.5 | 13.7 | 4.0 | 40.6 | 5 | 0.951 | 37.0 |
| A1 | 64 | 0.251 | 0.494 | 78.2 | 36.6 | 10.2 | 1.0 | 13.3 | 2.9 | 39.6 | 3.4 | 0.931 | 36.7 |
| A1 | 128 | 0.251 | 0.498 | 78.0 | 36.6 | 9.5 | 0.8 | 12.8 | 2.1 | 38.2 | 2.2 | 0.958 | 36.9 |
| A1 | 256 | 0.250 | 0.497 | 76.0 | 35.7 | 9.7 | 0.7 | 12.8 | 1.7 | 37.2 | 1.6 | 0.958 | 35.9 |
| A2 | 1 | 0.252 | 0.474 | 40.3 | 19.2 | 19.3 | 7.8 | 117.0 | 31.8 | 149.6 | 21.8 | 0.958 | 18.5 |
| A2 | 2 | 0.255 | 0.489 | 40.6 | 18.8 | 13.7 | 3.9 | 52.7 | 19.4 | 70.5 | 13.1 | 0.958 | 18.7 |
| A2 | 4 | 0.253 | 0.495 | 40.6 | 19.0 | 11.2 | 2.5 | 27.8 | 12.3 | 40.5 | 8.4 | 0.944 | 19.0 |
| A2 | 8 | 0.245 | 0.491 | 40.0 | 18.6 | 8.3 | 1.5 | 22.3 | 8.5 | 33.5 | 5.4 | 0.958 | 18.6 |
| A2 | 16 | 0.254 | 0.487 | 39.4 | 18.1 | 7.4 | 1.0 | 20.2 | 5.9 | 29.7 | 3.4 | 0.958 | 18.3 |
| A2 | 32 | 0.251 | 0.491 | 39.4 | 18.3 | 6.8 | 0.8 | 19.2 | 4.4 | 28.3 | 2.4 | 0.944 | 18.6 |
| A2 | 64 | 0.256 | 0.488 | 38.4 | 17.9 | 7.7 | 0.7 | 19.0 | 3.2 | 27.3 | 1.5 | 0.938 | 18.1 |
| A2 | 128 | 0.242 | 0.500 | 38.2 | 17.9 | 7.2 | 0.4 | 18.7 | 2.5 | 26.8 | 1.2 | 0.951 | 18.0 |
| A2 | 256 | 0.254 | 0.495 | 38.1 | 18.1 | 7.1 | 0.3 | 18.4 | 1.9 | 26.4 | 0.9 | 0.938 | 18.1 |
| A2b | 1 | 0.246 | 0.426 | 55.7 | 45.2 | 56.6 | 24.3 | 184.5 | 30.8 | 320.1 | 49.3 | 0.924 | 35.1 |
| A2b | 2 | 0.241 | 0.492 | 40.1 | 36.1 | 11.0 | 7.9 | 58.7 | 22.8 | 77.0 | 30.5 | 1.000 | 35.1 |
| A2b | 4 | 0.244 | 0.495 | 39.4 | 35.5 | 6.0 | 4.1 | 19.2 | 12.9 | 27.4 | 17.1 | 0.965 | 35.2 |
| A2b | 8 | 0.244 | 0.498 | 38.7 | 34.7 | 5.1 | 2.8 | 12.8 | 8.3 | 18.1 | 10.6 | 0.938 | 34.8 |
| A2b | 16 | 0.244 | 0.492 | 39.4 | 35.1 | 3.7 | 2.1 | 9.8 | 5.7 | 14.3 | 7.1 | 0.965 | 34.9 |
| A2b | 32 | 0.246 | 0.499 | 39.0 | 35.1 | 3.2 | 1.4 | 8.1 | 4.1 | 11.4 | 4.9 | 0.972 | 35.1 |
| A2b | 64 | 0.248 | 0.498 | 39.1 | 35.0 | 2.6 | 1.2 | 7.2 | 3.0 | 10.1 | 3.3 | 0.979 | 35.3 |
| A2b | 128 | 0.252 | 0.496 | 38.5 | 34.6 | 2.3 | 0.8 | 6.6 | 2.1 | 9.1 | 2.1 | 0.972 | 35.0 |
| A2b | 256 | 0.249 | 0.501 | 38.7 | 34.9 | 2.4 | 0.5 | 6.3 | 1.6 | 8.8 | 1.4 | 0.951 | 35.1 |

Table B1: Continued
| Scen | IDCat | $\hat{\lambda}_0$ | $\hat{\sigma}$ | $\hat{N}$ | $\hat{n}^{cap}$ | $\hat{N}$<br>RMSE | $\hat{n}^{cap}$<br>RMSE | $N$<br>CV | $\hat{n}^{cap}$<br>CV | $N$<br>Wid | $\hat{n}^{cap}$<br>Wid | $N$<br>Cov | $n^{cap}$ |
| --- | --- | --- | --- | --- | --- | --- | --- | --- | --- | --- | --- | --- | --- |
| A3 | 1 | 0.055 | 1.421 | 71.1 | 48.4 | 70.9 | 28.4 | 124.2 | 42.9 | 285.2 | 69.1 | 0.965 | 43.6 |
| A3 | 2 | 0.055 | 1.194 | 79.9 | 51.1 | 61.7 | 24.3 | 99.8 | 35.0 | 264.8 | 62.3 | 0.951 | 43.3 |
| A3 | 4 | 0.059 | 0.976 | 85.5 | 48.0 | 48.1 | 16.9 | 65.5 | 26.9 | 196.4 | 47.6 | 0.958 | 44.5 |
| A3 | 8 | 0.060 | 0.951 | 84.7 | 46.1 | 35.0 | 10.3 | 40.4 | 17.5 | 122.5 | 30.4 | 0.958 | 43.5 |
| A3 | 16 | 0.059 | 0.982 | 81.3 | 44.9 | 20.5 | 5.3 | 25.5 | 11.5 | 76.5 | 19.4 | 0.938 | 43.9 |
| A3 | 32 | 0.060 | 0.986 | 79.3 | 44.3 | 13.0 | 3.5 | 18.2 | 7.6 | 54.2 | 12.6 | 0.958 | 43.5 |
| A3 | 64 | 0.061 | 0.979 | 80.3 | 44.2 | 12.7 | 2.3 | 14.8 | 5.2 | 45.1 | 8.3 | 0.931 | 43.7 |
| A3 | 128 | 0.060 | 0.991 | 78.4 | 43.5 | 10.1 | 1.6 | 13.4 | 3.8 | 39.8 | 5.7 | 0.965 | 43.3 |
| A3 | 256 | 0.061 | 0.979 | 79.3 | 43.7 | 8.9 | 1.2 | 12.5 | 2.7 | 37.9 | 3.8 | 0.965 | 43.7 |
| A4 | 1 | 0.051 | 1.402 | 43.0 | 27.2 | 44.8 | 18.7 | 188.2 | 43.5 | 255.7 | 39.1 | 0.965 | 21.4 |
| A4 | 2 | 0.056 | 1.291 | 54.7 | 28.8 | 51.4 | 15.8 | 118.5 | 32.6 | 211.4 | 33.0 | 0.938 | 22.0 |
| A4 | 4 | 0.063 | 0.920 | 41.6 | 22.8 | 21.8 | 6.6 | 82.8 | 26.2 | 114.1 | 22.0 | 0.972 | 21.4 |
| A4 | 8 | 0.058 | 0.964 | 41.8 | 23.0 | 16.3 | 4.2 | 48.7 | 17.5 | 70.0 | 14.9 | 0.958 | 22.2 |
| A4 | 16 | 0.060 | 0.975 | 40.0 | 22.1 | 11.3 | 2.8 | 29.9 | 11.4 | 42.9 | 9.1 | 0.924 | 21.8 |
| A4 | 32 | 0.061 | 0.962 | 40.4 | 22.0 | 9.2 | 2.0 | 23.3 | 7.8 | 34.6 | 5.9 | 0.938 | 22.0 |
| A4 | 64 | 0.060 | 0.989 | 39.0 | 21.6 | 7.2 | 1.3 | 20.1 | 5.7 | 29.0 | 4.0 | 0.958 | 21.7 |
| A4 | 128 | 0.062 | 0.970 | 40.0 | 22.0 | 7.2 | 0.9 | 18.4 | 4.1 | 27.5 | 2.6 | 0.944 | 22.1 |
| A4 | 256 | 0.063 | 0.969 | 39.0 | 21.6 | 6.8 | 0.6 | 17.6 | 3.0 | 25.6 | 1.7 | 0.951 | 21.7 |

## 10 Supplement 1: MCMC mixing

Here, we demonstrate some features of the mixing of categorical SPIM MCMC chains. First, as more loci or identity categories are added, the uncertainty in *n^cap^*, the number of unique individuals captured is reduced, and this in turn can lead to very low probability values of *n^cap^* that may require many MCMC iterations to fully characterize the posterior of *N*. This can be seen in Figures S1 and S2, depicting 32 MCMC chains for *n^cap^* for the 1-7 loci data sets of Kentucky male and female bears, respectively. These figures show that as the genotype information becomes more rich, *n^cap^* is increasingly constrained, and the largest values with positive posterior probability become increasingly less probable, requiring longer chains to record the increasingly rare events that *n^cap^* takes these larger values. Specifically, for the female data set, *n^cap^* = 46 becomes increasingly less probable and for the male data set, *n^cap^* = 38 becomes increasingly less probable (true values are 45 and 37, respectively).

Figure S3 depicting a single MCMC chain from the 6 loci male data set shows that these very small shifts in *n^cap^* can have a non-negligible impact on the estimates of the other parameter values which can lead to bimodal posteriors for *n^cap^*, *λ_0_*, *σ*, and *N*. This pattern was more pronounced in the male data set due to an especially long-distance spatial recapture for a single individual. When *n^cap^* was estimated to be 38, this individual’s captures were split across two latent individuals, while when *n^cap^* was estimated to be 37, this individual’s captures were correctly combined into a single latent individual. However, due to the distance between this individual’s captures, the estimate of *σ* had to shift upwards substantially to make this combination of samples more likely and the shift in *σ* propagated to *λ*_0_ and *N*. Note also, that the *n^cap^* posteriors for male data sets with higher numbers of loci tend to stay at these low probability *n^cap^* values for many consecutive iterations, while the *n^cap^* posteriors for the female data sets do not remain at the low probability *n^cap^* values for many consecutive iterations. This can be attributed to the bimodal distribution of *σ* in the male data set-when *n^cap^* = 38, *σ* is underestimated relative to when *n^cap^* = 37, leading to an exceeding low probability that the samples from the individual with the long distance spatial recapture will be proposed to be combined into a single latent individual.

We suspect bimodal posteriors (possibly multimodal) will be more common in two scenarios. First, when *n_cap_* is small, a 1 unit difference in *n_cap_* should have a larger influence on the estimate of *N*. Second, realizations of the capture process leading to one or a few long distance spatial recaptures relative to *σ* as seen in the male data set may often lead to bimodal posteriors. These long distance spatial recaptures could be due to rare realizations of the capture process with fixed detection function parameters, but will be more common if there is individual heterogeneity in detection function parameters, especially *σ*. If individual heterogeneity in *σ* explains the long distance spatial recapture in the male data set, the 1-7 loci analyses suggest that the misspecification of the observation process that leads to a very low probability this individual’s samples all came from the same latent individual can be overcome by adding more genotype information, making it increasingly unlikely that two individuals in the population had the same genotype.

Due to the possibility of bimodal posteriors with low probability modes that may have a disproportionate impact on the posterior, we suggest running multiple chains for many iterations when there is little to no apparent uncertainty in *n_cap_*.

**Figure S1:**
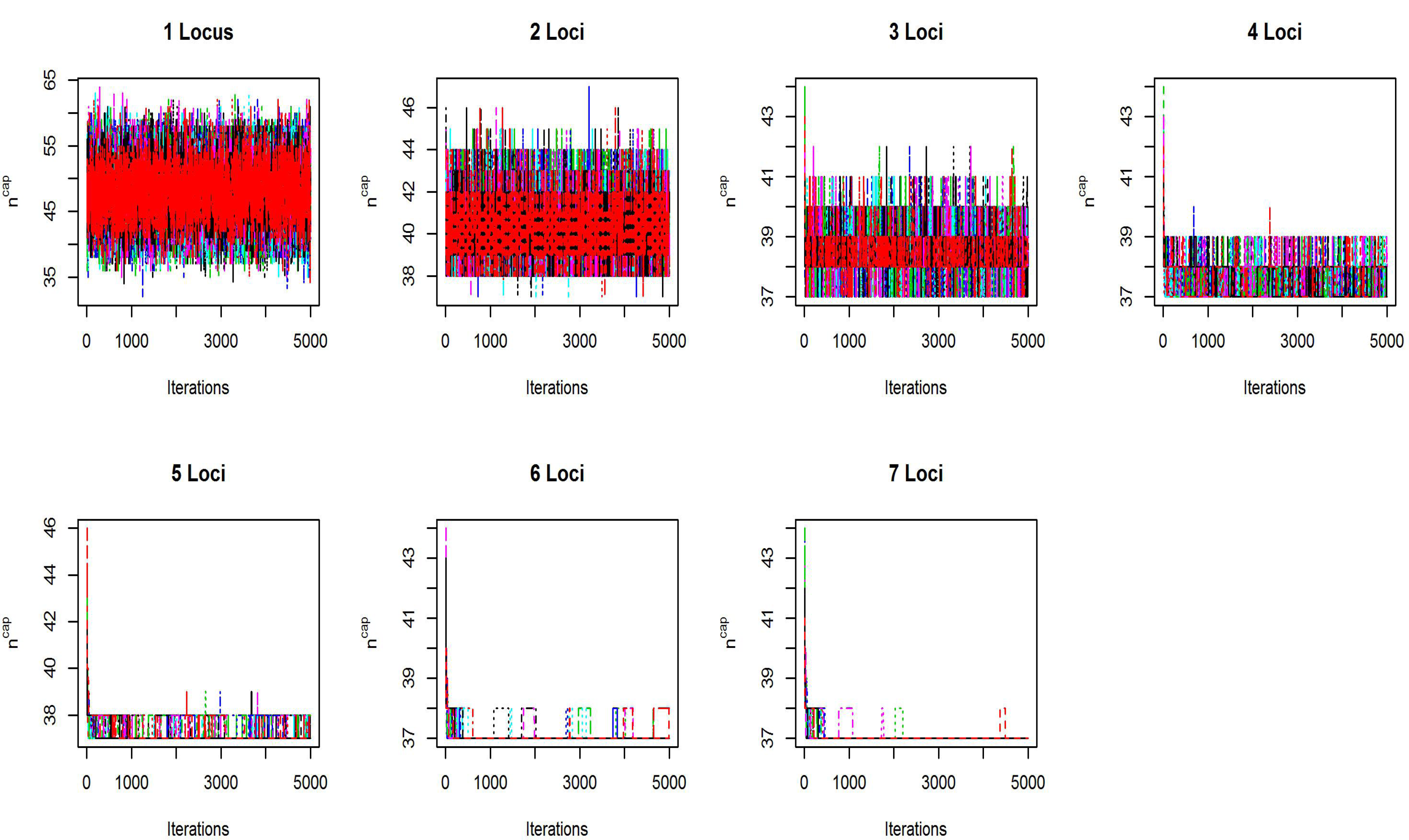
MCMC trace plots for *n^cap^*, the number of individuals captured, for the 1-7 loci KY male bear data sets. 32 chains were run for 250, 0 iterations, thinned by 50. Burn in samples were not removed in these plots to show convergence.

**Figure S2:**
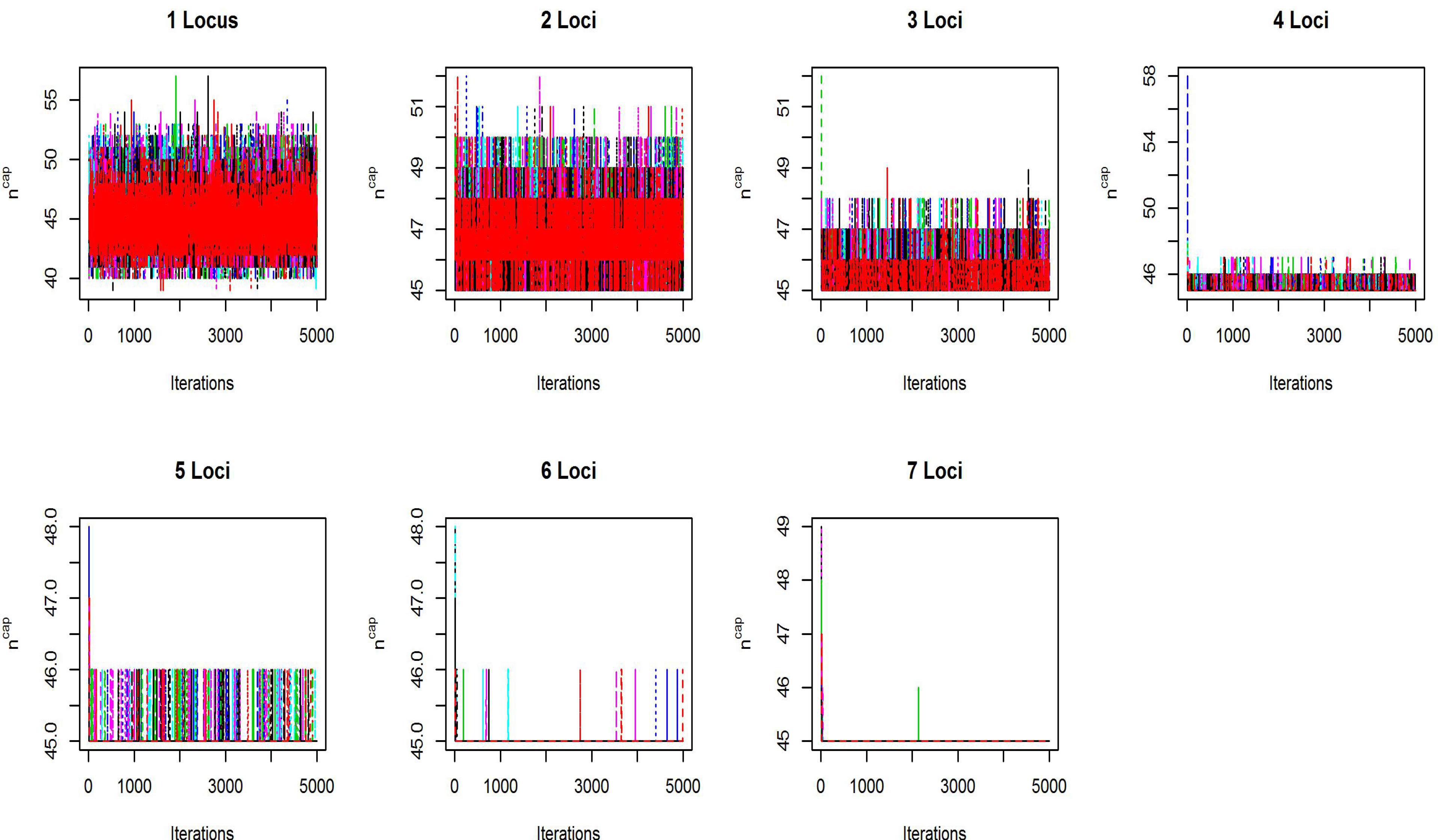
MCMC trace plots for *n^cap^*, the number of individuals captured, for the 1-7 loci KY female bear data sets. 32 chains were run for 250,000 iterations, thinned by 50. Burn in samples were not removed in these plots to show convergence.

**Figure S3:**
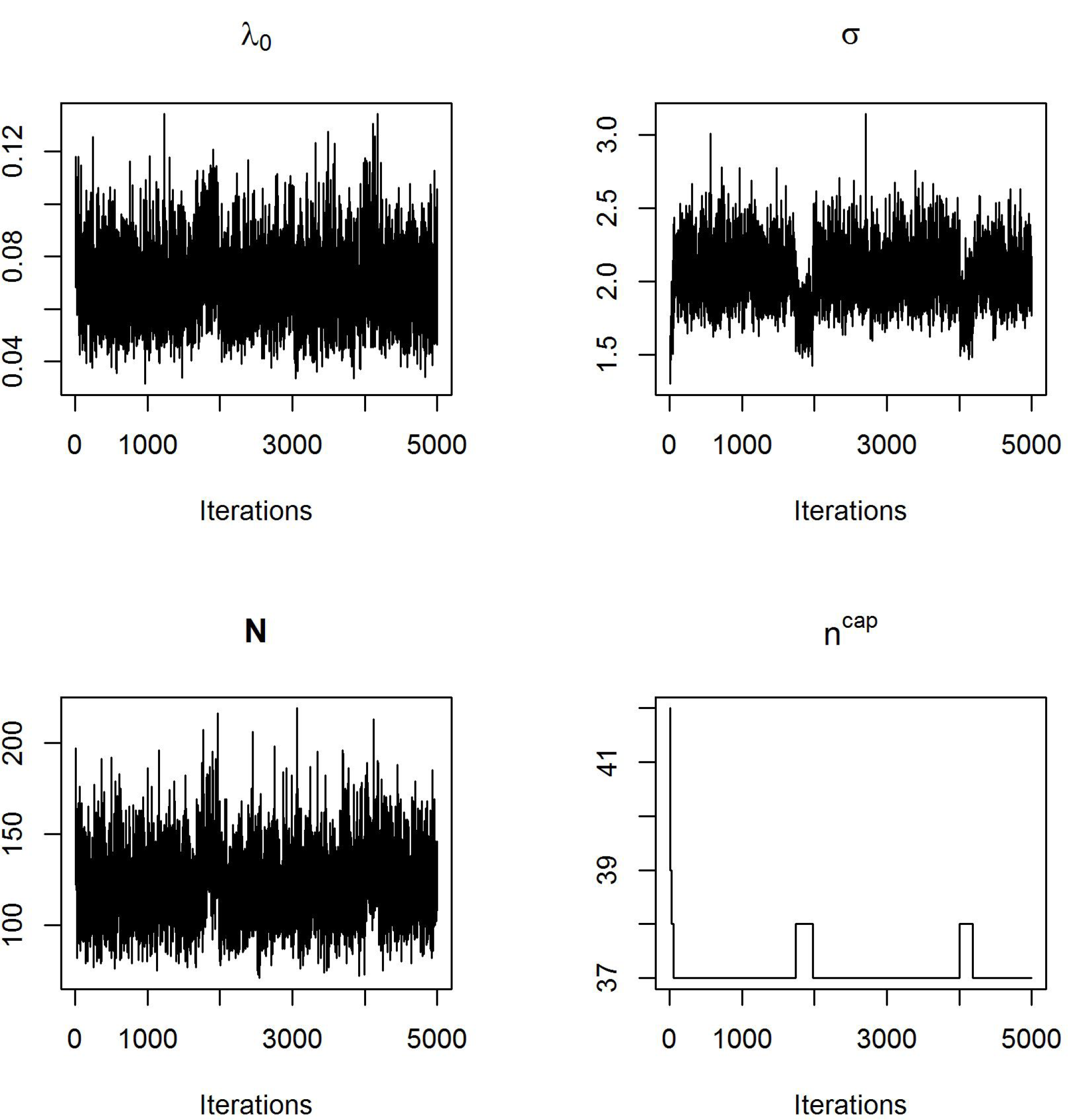
An example MCMC trace plot of a single MCMC chain from the 6 loci KY male bear data set exhibiting bimodal posterior distributions. Two upward shifts in *n^cap^*, cause downward shifts in σ, which propagate to, λ_0_ and *N*.

